# A macrocyclic peptide-based fusion inhibitor targeting SARS-CoV-2 Spike S2 subunit

**DOI:** 10.64898/2026.03.04.703990

**Authors:** Hirofumi Ohashi, Tatsuro Kawamura, Masaki Ohuchi, Haruaki Kurasaki, Naoko Iwata-Yoshikawa, Yuichiro Hirata, Saya Moriyama, Kaho Shionoya, Kazutaka Nagatomo, Takayuki Nagasawa, Junpei Yamamoto, Kei Sudo, Naoko Nakamura, Katsuma Matsui, Haruhiko Ogawa, Kouji Yoshida, Yoshiaki Shimada, Tomoko Maruyama, Tomoaki Higuchi, Shoko Ito, Yoshimasa Takahashi, Naoki Kawamura, Patrick C. Reid, Masato Murakami, Tadaki Suzuki, Noriyo Nagata, Hidetomo Kitamura, Koichi Watashi

## Abstract

Continuous emergence of SARS-CoV-2 variants carrying mutations in Spike presents a significant challenge for durable antiviral agents. Here we screen for random 13-amino acid non-mimetic macrocyclic peptides that bind to Spike and identify PA-001 that inhibits SARS-CoV-2 infection with high potency at 0.23–2.9 nM as 50% inhibitory concentration (IC_50_). PA-001 bound to Spike S2 subunit and inhibited the membrane fusion during virus entry. Through drug-resistant selection, we revealed that PA-001 targeted the fusion peptide proximal region (FPPR) in S2, which has not been recognized as a drug target to date. Consistent with its highly conserved amino acid sequences beyond strains, PA-001 exhibited broad antiviral activity against all tested SARS-CoV-2 variants, in contrast to clinically-approved S1-targeting antibodies that lost activity to Omicron variants. PA-001 suppressed SARS-CoV-2 propagation and disease progression in mouse- and hamster-infection models, both by administration prophylactically and therapeutically. Combination therapy with remdesivir further enhanced antiviral profiles. In clinical phase-I trial, PA-001 was well-tolerated and showed high systemic exposure, with 4,300–10,300-fold concentration of IC_50_ as maximum plasma concentration by single administration to healthy volunteers. These evidence propose FPPR as an unexpected antiviral drug target accessible by macrocyclic peptides and identify PA-001 as a potent anti-SARS-CoV-2 fusion inhibitor.

## Introduction

Coronavirus disease 2019 (COVID-19), caused by infection with severe acute respiratory syndrome coronavirus 2 (SARS-CoV-2), has had a profound impact on global public health, resulting in more than 770 million cumulative cases and over 7 million deaths worldwide (https://covid19.who.int/). Despite the development and widespread deployment of vaccines and antiviral drugs, COVID-19 continues to impose a substantial disease burden, especially among people including older adults, individuals with underlying medical conditions, and immunocompromised populations^1-3^. Throughout and beyond the pandemic period, SARS-CoV-2 has exhibited continuous viral evolution, given rise to numerous variants characterized by the accumulated mutations^4^. These genetic changes have altered viral pathogenicity and transmissibility and, critically, reduced susceptibility to approved antiviral drugs. There is a need to develop novel antiviral agents sustainably retaining their efficacy against ongoing emergence of SARS-CoV-2 variants.

The Spike glycoprotein of SARS-CoV-2 mediates viral entry to host cells through receptor binding and membrane fusion. Spike constitutes a trimeric precursor and comprises two functional subunits, S1 and S2. S1 consists of N-terminal domain (NTD), receptor binding domain (RBD), and C-terminal domains (CTDs), and is essential for recognition of host entry receptor, angiotensin converting enzyme 2 (ACE2), through its RBD^5,6^. S2 contains multiple functional domains including fusion peptide (FP) and its proximal region (FPPR), heptad repeat 1 (HR1), central helix (CH), connector domain (CD), heptad repeat 2 (HR2), and transmembrane domain (TMD) (see Fig. 2E)^4,6^. Upon RBD in the S1 subunit engagement to ACE2, S2 is processed at its N-terminal S2′ site by host proteases, either transmembrane serine protease 2 (TMPRSS2) on the cell surface or cathepsin L in the endosome, which then induces large-scale conformational rearrangements of S2 in the membrane fusion process (Supplementary Fig. 1)^6-12^: The S1-ACE2 complex dissociates from the S2 trimer (Supplementary Fig. 1a-c), and the original conformation (pre-fusion form) of S2 is converted to forms a long central coiled coil with blunted cone-shaped structure (involving FP and the adjacent FPPR) at its top, which allows insertion into the target cell membrane (Intermediation form; Supplementary Fig. 1d). It subsequently refolds into the stable structure by interaction of HR1 and HR2 to form a six-helix bundle that constitutes a rigid, rod-like shape (Post-fusion form; Supplementary Fig. 1e-f) to undergo fusion of viral envelope to cellular membranes.

This viral entry process is an attractive antiviral target^13^. The clinically-approved antibodies for COVID-19 treatment, such as casirivimab, imdevimab, sotrovimab, tixagevimab, cilgavimab, and bebtelovimab, recognize the S1 subunit and are designed to strongly block the S1 RBD-ACE2 binding^14,15^. However, their neutralization activities have been lost by accumulated mutations, especially in RBD, in the recent Omicron subvariants such as BQ.1 and XBB^16^. Other S1-targeting antibodies, including those targeting NTD, also face the similar challenges for emerging escape variants due to high sequence variability in this region. Given its highly conserved sequence among circulating SARS-CoV-2 variants, S2 is expected as an alternative target for broadly active antivirals. So far, S2-targeting antibodies have been identified, including those recognizing the region downstream of the S2′ cleavage site (used to be referred as FP, but is presently defined an intermediate loop) and the stem helix at the upstream of HR2^17-19^, but their neutralizing activities were typically low compared to those targeting RBD in S1. So far, peptides mimicking the HR2 sequence aiming to inhibit HR1-HR2 interaction during the conversion to the post-fusion form have been under clinical development. These peptides have shown broad anti-SARS-CoV-2 activity, but their original activities of pre-optimized peptides were 500-3,000 nM ranges as 50% inhibitory concentrations (IC_50_)^18,20^. Later, these S2-mimicking linear peptides have been optimized for improving their activities through polyethylene glycol or cholesterol conjugations. These antibody- and linear peptide-based approaches face a challenge in which their target site as well as antiviral activities are limited due to the S2’s inherent characteristics including limited surface area accessibility by large molecules as well as low immunogenicity and high flexibility.

Cyclic peptide-based drugs are expected to provide an alternative approach^21^ that can have drug-like properties and advantages including accessibility to narrow target sites, high specificity, tunable stability and efficacy, low immunogenicity, and relatively less cost compared to antibodies. In this study, we show that a non-mimetic macrocyclic peptide, PA-001, inhibits infection of broad SARS-CoV-2 variants via targeting FPPR within the S2 subunit, which is highly conserved in sequence but previously undescribed as a druggable site. By using SARS-CoV-2-inoculated mouse and hamster models, PA-001 suppressed viral propagation and ameliorated disease progression under administrations either prophylactically or therapeutically. A first-in-human Phase I dose-escalation trial demonstrated a favorable safety profile with high systemic exposure in pharmacokinetics (achieving maximum plasma concentration of over 4,000-fold concentration of in vitro IC_50_ against Omicron strains) in healthy volunteers. Our findings present PA-001 as an anti-SARS-CoV-2 drug candidate and that FPPR is a potential druggable target for SARS-CoV-2 fusion inhibitor.

## Results

### Identification of an anti-SARS-CoV-2 cyclic peptide, PA-001

To explore novel COVID-19 therapy targeting SARS-CoV-2 Spike, we discovered and developed anti-SARS-CoV-2 agents following the scheme as summarized in Fig. 1A. Through screening of a library consisting of ∼10^13^ random peptides using Peptide Discovery Platform System (PDPS), which we have established so far^22-24^ (Fig. 1B), and the following rational optimization process through medicinal-chemistry program, we developed PA-001 (Fig. 1C) as an investigational agent currently in clinical development as a therapy against COVID-19. In the present study, we report the in vitro and in vivo antiviral profile of PA-001.

**Fig. 1.**
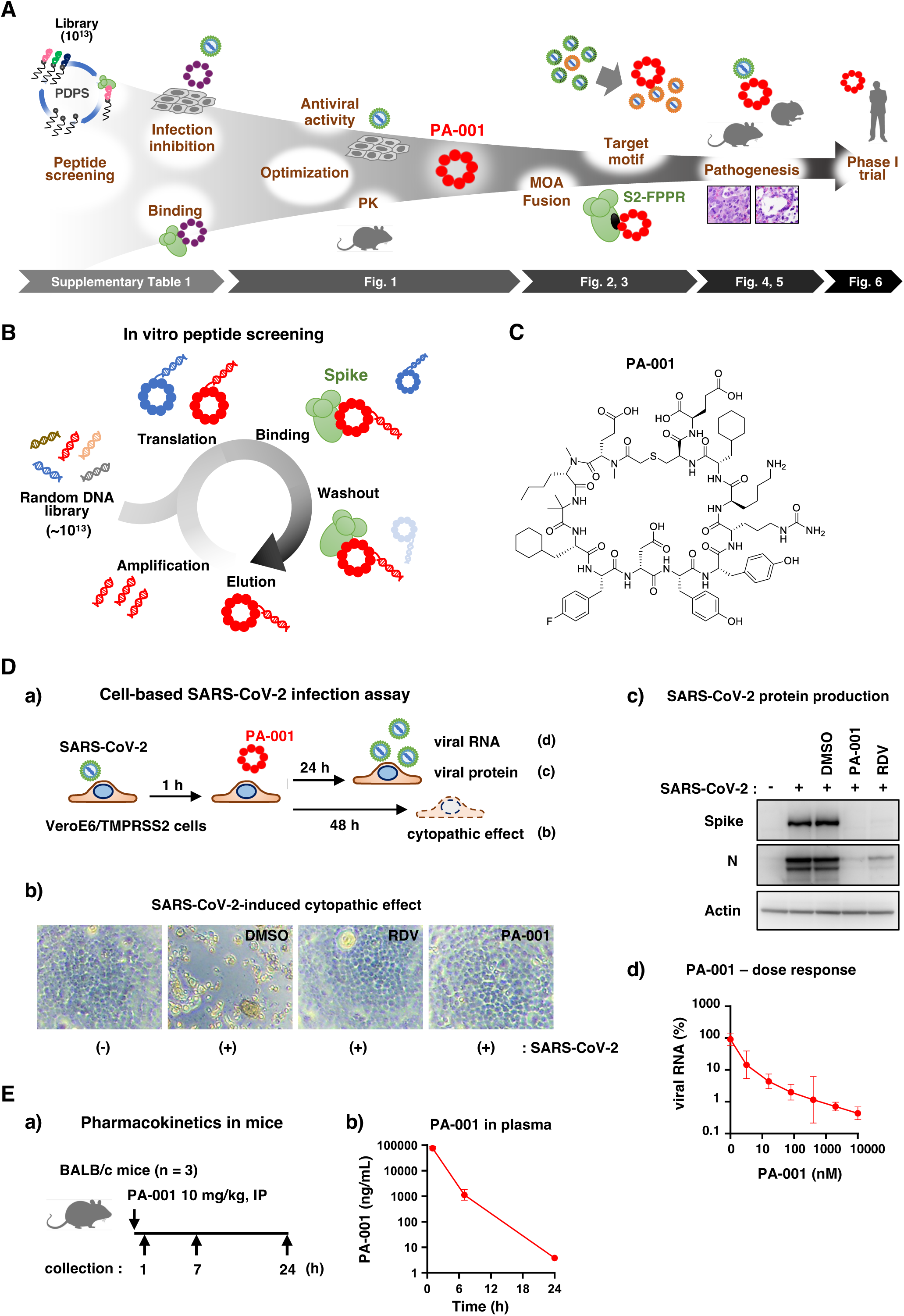
Identification of PA-001, a cyclic peptide showing potent anti-SARS-CoV-2 activity. (A) Schematic diagram of the flow to identify Spike-binding macrocyclic peptides and evaluate their antiviral activity. The bottom arrows show the figures corresponding to the analyses shown above. (B) Schematic representation of the mRNA display system to screen for Spike-binding macrocyclic peptides. (C) Structure of PA-001. (D) PA-001 inhibited the propagation of SARS-CoV-2. (a) Schematic representation of SARS-CoV-2 infection assay. VeroE6/TMPRSS2 cells were inoculated with or without SARS-CoV-2 (Wk-521; wuhan strain) at an MOI of 0.003 for 1 h, followed by washing out free virus and culturing cells with the medium supplemented with indicated compounds for additional 24 h to detect viral proteins in cells by immunoblot (c) and to quantify viral RNA in the culture supernatant by real time RT-PCR (d), as well as culturing cells for 48 h to observe virus-induced cytopathic effect (CPE) by microscopic observation (b). In (c), viral Spike (upper) and N (middle) proteins as well as cellular actin as an internal control (lower) were detected. In (d), quantified viral RNAs were plotted against the concentration of treated PA-001 (nM). (E) Pharmacokinetics of PA-001 in mice. (a) Scheme for treatment with PA-001 BALB/c mice and time points (1, 7, and 24 h post IP injection) of plasma collection. (b) BALB/c mice were intraperitoneally administered with PA-001 at 10 mg/kg and were chased to obtain plasma at 1, 7, and 24 h post-administration to quantify PA-001 concentration (mean±SD).

Peptides that were identified to bind Spike in vitro were optimized to improve antiviral potency and pharmacokinetics (Fig. 1A). We examined i) affinity with Spike in vitro (Supplementary Table 1), ii) inhibition activity to SARS-CoV-2 infection in a cell-based assay (Fig. 1D), and iii) pharmacokinetics profiles in mice (Fig. 1E). In surface plasmon resonance, PA-001 had high affinity to recombinant Spike, with 1.3 nM as a K_D_ (Supplementary Table 1). Activity of peptides to inhibit SARS-CoV-2 infection was examined in a cell-based SARS-CoV-2 infection assay: VeroE6/TMPRSS2 cells^25^ were inoculated with a SARS-CoV-2 clinical isolate (Wk-5.2.1, Wuhan type) at a multiplicity of infection (MOI) of 0.003 for 1 h and were cultured in the presence or absence of peptides for 48 h to assess SARS-CoV-2-induced cytopathology (Fig. 1D). The cell line exhibited robust cytopathology by SARS-CoV-2 propagation at 48 h after infection, which was however completely blocked by treatment with 50 nM PA-001, as 10 μM remdesivir (RDV), a clinically-approved anti-SARS-CoV-2 polymerase inhibitor^26^ used as a positive control (Fig. 1D-b). PA-001 reduced the production of SARS-CoV-2 proteins, Spike and N as representatives detected by immunoblotting, at 24 h post-inoculation, the time before showing cytopathic effect (Fig. 1D-c). PA-001 reduced viral RNA at 24 h post-inoculation in a dose-dependent manner (Fig. 1D-d). Pharmacokinetics of a single intraperitoneal administration with PA-001 at 10 mg/kg to BALB/c mice showed 76,667 ng/mL (41,500 nM) as maximum plasma concentration (C_max_) and a gradual decline but sustained plasma concentration, with 1.0 h as half-life, and 282,236 ng·h/mL (153,000 nM·h) as area under the curve (AUC_0-24_) in plasma (Fig. 1E, Supplementary Table 2). From these data showing the strong potential of PA-001 as an anti-SARS-CoV-2 agent, we focused on PA-001 in the following analysis.

### PA-001 targets S2 and inhibits Spike-dependent membrane fusion

We determined the target of PA-001 by in vitro binding assay that incubated PA-001 at varying concentrations with recombinant either full-length (S1+S2), S1, or S2 protein as targets. As shown in Fig. 2A, PA-001 presented the binding signal with the full-length Spike and S2 in a dose-dependent manner, but not with S1 at any treated concentrations up to 100 nM (Fig. 2A), suggesting that PA-001 binds to S2. In the SARS-CoV-2 life cycle, cell entry is initiated by viral attachment to cells through binding of S1 RBD to ACE2 on cell surface (Fig. 2B, red); This is followed by the fusion of viral envelope with cellular membranes either on cell surface or at intracellular vesicles, triggered by cleavage of the S2 internal site (S2′ site) by cellular proteases (Fig. 2B, brown). To confirm the target step of PA-001 during the entry process, we separately assessed the activity of viral attachment to cell surface (Fig. 2C-a) and membrane fusion (Fig. 2C-b). Attachment assay was performed by incubating VeroE6/TMPRSS2 cells with SARS-CoV-2 at 4°C to allow viral-cell attachment but not the following processes, followed by quantification of viral RNA on the cell surface. Cell-attached SARS-CoV-2 RNA was dose-dependently reduced upon treatment with casirivimab + imdevimab (CAS + IMD) cocktail antibodies, clinically-approved SARS-CoV-2 neutralizing antibodies^27^ used as a positive control, whereas that was not significantly affected by treatment with PA-001, as with mefloquine, a reported post-attachment entry inhibitor^28^ used as a negative control (Fig. 2C-a). On the other hand, in the cell fusion assay, which coincubated Spike-overexpressing HEK293T cells with endogenously ACE2-expressing VeroE6/TMPRSS2 cells to induce cell membrane fusion, essentially as previously described^29^, PA-001 clearly reduced Spike-ACE2-induced fusion in a dose-dependent manner, and its IC_50_ was calculated to be 3.6 nM (Fig. 2C-b). These results are consistent with that PA-001 targets S2, which mediates the membrane fusion process.

**Fig. 2.**
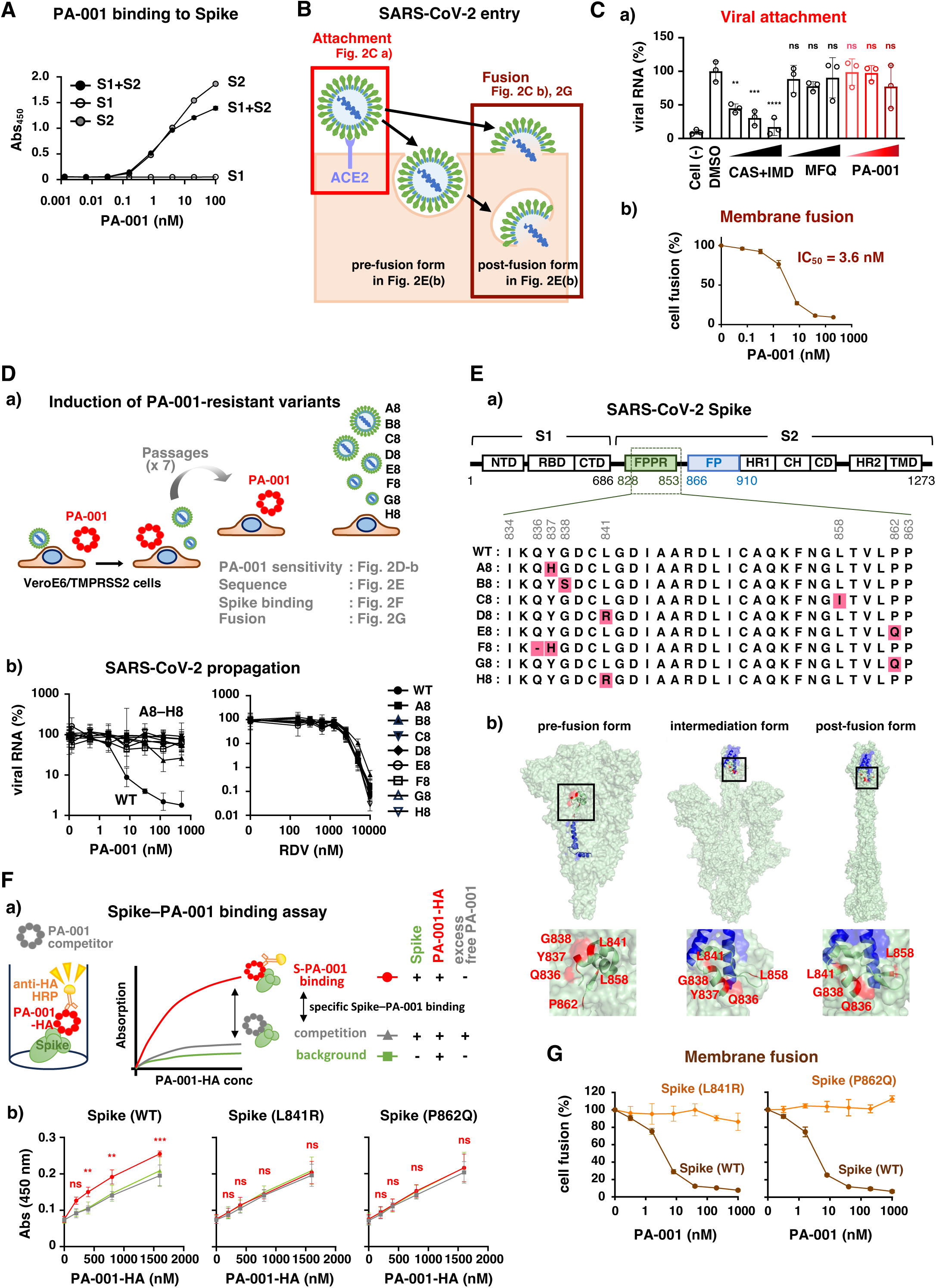
PA-001 bound to FPPR in the S2 subunit and inhibited the membrane fusion. (A) Binding of PA-001 to recombinant SARS-CoV-2 Spike protein of the full length (S1+S2), S1, or S2 regions. Each recombinant Spike protein was immobilized and incubated with HA-tagged PA-001 at various concentrations up to 100 nM, followed by washing out and detection with horseradish peroxidase (HRP)-conjugated anti-HA antibody. (B) Schematic model for the SARS-CoV-2 entry process, in which viral attachment to cells via Spike-ACE2 binding is followed by the fusion of viral envelope with cellular membrane either on the cell surface or in the endosome/lysosome. (C) (a) Virus-cell attachment assay. VeroE6/TMPRSS2 cells were exposed to SARS-CoV-2 at an MOI of 0.01 at 4°C for 15 min in the presence or absence of casirivimab (CAS) + imdevimab (IMD) (50, 500, or 5,000 ng/mL), mefloquine (MFQ) (0.4, 2, or 10 μM), or PA-001 (40, 200, or 1,000 nM), and were then washed out to quantify cell-bound viral RNA by real-time RT-PCR. Ordinary one-way ANOVA followed by Dunnett’s multiple comparisons with the DMSO control was performed (*: p<0.05, **: p<0.01, ***: p<0.001, ****: p<0.0001). (b) Cell-cell fusion assay. Spike-overexpressing HEK293T cells carrying the LgBiT gene were co-cultured with ACE2-expressing VeroE6/TMPRSS2 cells carrying the HiBiT gene upon treatment with or without varying concentrations (up to 200 nM) of PA-001 for 1 h to detect luciferase activity to evaluate fusion activity mediated by Spike and ACE2. IC_50_ of PA-001 is also indicated. (D) Induction of PA-001-resistant SARS-CoV-2 variants. (a) Schematic representation of the infection assay for inducing PA-001-resistant SARS-CoV-2 clones. SARS-CoV-2-infected VeroE6/TMPRSS2 cells were continuously cultured with PA-001 at 1,000 nM for 24-48 h (Passage 0). The resultant culture supernatants were collected and diluted to 10-fold, which were then inoculated to naive VeroE6/TMPRSS2 cells in the presence of 1,000 nM PA-001 and cultured for 24 h (Passage 1). The culture supernatants were repeatedly collected and then inoculated to the cells upon 1,000 nM PA-001 for the next passage. At passage 7, the culture supernatants were extracted from eight independent cultures (A8 – H8). (b) PA-001 sensitivity of the collected supernatant clones. VeroE6/TMPRSS2 cells were inoculated with these collected supernatants or the parental virus (WT) upon treatment with varying concentrations of PA-001 (left) or RDV as a reference (right) to determine virus RNA levels in the culture supernatants as shown in Fig. 1D-d. (E) Amino acid sequence of the PA-001-resistant clones. (a) Upper, schematic structure for the SARS-CoV-2 Spike protein. Lower, amino acid sequence of the clones, A8–H8, as well as the parental virus (WT). The substituted amino acids are highlighted by red. The amino acid numbers are indicated. (b) Structure of Spike trimer in the pre-fusion (left, PDB: 6XR8), intermediation (center, PDB: 8Z7P), and post-fusion (right, PDB: 8FDW) forms. Red color indicates the mutated amino acids shown in (a). Blue color indicates FP. (F) Binding capacity of recombinant SARS-CoV-2 Spike carrying the L841R and P862Q substitutions to PA-001. (a) Each recombinant Spike (WT, L841R, and P862Q) was immobilized and incubated with HA-tagged PA-001 (PA-001-HA) at varying concentrations in the presence (gray, competition) or absence (red, S-PA001 binding) of non-tagged PA-001 at 10 μM. After washing out, bound HA-PA-001 was visualized with HRP-conjugated anti-HA antibody. Samples without Spike-immobilization were also prepared to define the background signal (green). (b) Binding signals with Spike WT (left), Spike L841R (center), and Spike P862Q (right) are plotted against the concentrations of treated PA-001-HA. Ordinary two-way ANOVA followed by tukey’s multiple comparisons was performed (*: p<0.05, **: p<0.01, ***: p<0.001). (G) Cell fusion assay performed as described in Fig. 2C-b using the cells overexpressing either Spike WT, L841R, or P862Q co-cultured with ACE2-expressing cells upon PA-001 treatment at various concentrations.

To identify the S2 target site responsible for the PA-001 sensitivity, we isolated PA-001-resistant SARS-CoV-2 variants by continuous exposure of SARS-CoV-2-infected cells with PA-001 and serial passages, essentially described previously^30^ (Fig. 2D-a): VeroE6/TMPRSS2 cells infected with SARS-CoV-2 were continuously treated with PA-001 and their culture supernatant was reinfected to naive VeroE6/TMPRSS2 cells. After seven passages, we collected SARS-CoV-2 in the culture supernatant from eight independent cell batches (A8, B8, C8, D8, E8, F8, G8, and H8). We then examined the sensitivity of these virus clones to PA-001, as well as RDV as a reference, by inoculating the cells with these viruses or the parental wild type SARS-CoV-2 and treated with PA-001 or RDV at varying concentrations to quantify viral RNA produced into the culture supernatant. As shown in Fig. 2D-b, all the virus clones (A8-H8) showed no or limited reduction in viral RNA levels by treatment with PA-001 up to 500 nM, in contrast to the drastic reduction in RNA of the parental SARS-CoV-2 (Fig. 2D-b left, and Supplementary Fig. 2A-a). In contrast, all these variants remained sensitive to RDV, as the parental SARS-CoV-2 (Fig. 2D-b right, Supplementary Fig. 2A-b). The calculated 90% inhibitory concentrations (IC_90_s) to PA-001 for all the clones were >500 nM (v.s. 8.4 nM for the parental virus), whereas those to RDV for all clones remained 3,000-4,500 nM that were equivalent to the parental virus (Supplementary Fig. 2B), indicating that all the isolated variants acquired specific resistance to PA-001.

Next-generation sequencing of viral RNA derived from these eight clones identified 1-3 amino acid substitutions for each clone, with enriched substitutions in the S2 region (Supplementary Fig. 2C), consistent with that PA-001 targets S2. Interestingly, D8 and H8 had the identical and single mutation that caused amino acid substitution, L841R in S2 protein (Fig. 2E-a, Supplementary Fig. 2C), suggesting that the L841R substitution is responsible for acquiring the PA-001 resistance. Similarly, E8 and G8 possessed the identical but another single amino acid substitution, P862Q in S2 (Fig. 2E-a, Supplementary Fig. 2C). Remarkably, all the remaining clones also acquired a substitution cluster at the neighboring amino acids, Q836 (in F8), Y837 (in A8 and F8), G838 (in B8), and L858 (in C8) (Fig. 2E-a highlighted by pink, Supplementary Fig. 2C). All these amino acids reside at the upstream of the fusion peptide (FP: 866-910 aa) in S2, referred as the fusion peptide proximal region (FPPR)^4,6-9^ (Fig. 2E-a). Based on the reported three-dimensional structure of Spike protein at pre-fusion, intermediation, and post-fusion forms^10-12^, FPPR dynamically moves and changes its structure, together with FP, during the conversion from the pre-fusion to the post-fusion form^4^: FPPR is originally exposed on the surface in a concave area of the central part of the pre-fusion trimer, and undergoes a dynamic locational change to the top of S2, forming the cone-shaped structure to assist FP insertion into target membrane in the post-fusion form^11^ (Fig. 2E-b).

As the above critical amino acids, Q836, Y837, G838, L841, L858, and P862 are exposed on the surface of the Spike pre-fusion trimer, we assume these amino acids constitute the interface to PA-001 binding, of which amino acid substitutions lead to escape from PA-001 binding. We then examined the binding activity of recombinant Spike having a substitution of either L841R (in D8 and H8) or P862Q (in E8 and G8) to PA-001 by ELISA: Spike proteins were immobilized on the well, and incubated with varying amount of HA-tagged PA-001, which was then visualized by adding anti-HA-conjugated horseradish peroxidase (HRP). Specific Spike-PA-001 binding was evaluated by the difference in the signal from that under coincubation with and without excess amount of non-HA-tagged PA-001 as a competitor (Fig. 2F-a). This specific Spike-PA-001 binding was observed along with the increasing amount of PA-001-HA by using the wild type Spike, however, this signal completely disappeared for Spike carrying the substitution at either L841R or P862Q (Fig. 2F-b), suggesting that these amino acid substitutions abrogate PA-001 binding. Consistently, PA-001’s dose-dependent inhibition of the Spike-ACE2-mediated cell membrane fusion was totally cancelled up to 1,000 nM by mutations of L841R or P862Q in Spike (Fig. 2G). These data cumulatively suggest that PA-001 binds to FPPR to inhibit S2-dependent membrane fusion.

### PA-001 shows antiviral activity to broad range of SARS-CoV-2 variants

Multiple neutralizing antibodies have been approved for clinical use, however, all of them have lost antiviral efficacy by amino acid substitutions in Spike observed in naturally-occurring SARS-CoV-2 variants^14,31^. Most of these substitutions were concentrated in the S1 subunit. To clarify the breadth of the antiviral activity of PA-001, we infected VeroE6/TMPRSS2 cells with SARS-CoV-2 Wuhan, Alpha, Beta, Gamma, Delta, and Omicron (BA.1.18 and XBB.1.16), and treated with varying concentrations of PA-001 as well as CAS + IMD as a reference to quantify viral RNA in the culture supernatant. As previously reported, CAS + IMD showed antiviral effects on Wuhan to Delta variants, but not Omicron variants (Fig. 3A, lower graphs), because of the amino acid substitutions in RBD of S1^32,33^. In contrast, PA-001 reduced SARS-CoV-2 infection for all the variants examined including Omicron variants, BA.1.18 and XBB.1.16, in a dose-dependent manner (Fig. 3A, upper graphs). Anti-SARS-CoV-2 activities of PA-001 against all these variants ranged 0.23-2.9 nM as IC_50_s and 1.3-9.4 nM as IC_90_s, in contrast to the complete loss of activity of CAS + IMD against BA.1.18 and XBB.1.16 (IC_50_ and IC_90_ >1,000 ng/mL) (Supplementary Table 3). Furthermore, PA-001 showed the inhibitory activity to the recent Omicron subvariants, JN.1 and XFG, in contrast to loss of activity of CAS + IMD, examined by infection of VSV-based pseudovirus carrying Spike derived from each SARS-CoV-2 variant, as previously described^34,35^ (Supplementary Fig. 3A). Consistently, the amino acid sequences of all these variants from 834 to 863 aa are almost completely conserved, except for a single non-essential substitution at position 856 only in BA.1.18 (Fig. 3B, Supplementary Fig. 3B). Thus, PA-001 showed an advantage that was effective on broad range of naturally-occurring SARS-CoV-2 variants, in clear contrast to the clinically-approved anti-S1 antibodies.

**Fig. 3.**
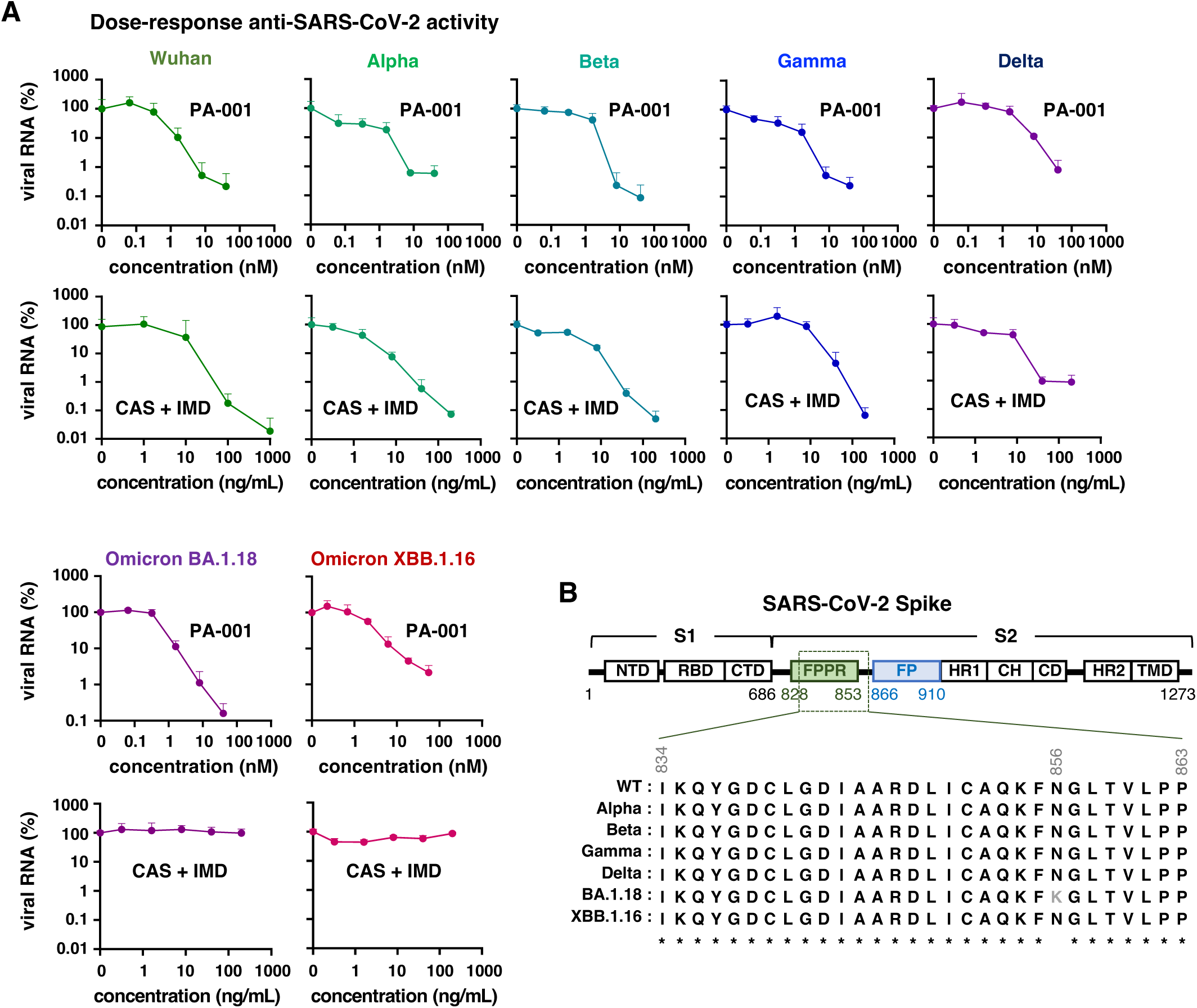
PA-001 showed a broad anti-SARS-CoV-2 activity against multiple variants. (A) Antiviral activity of PA-001 and casirivimab + imdevimab against the SARS-CoV-2 variants (Wuhan, Alpha, Beta, Gamma, Delta, and Omicrons BA.1.18 and XBB.1.16). Drug response was evaluated as described in Fig. 1D-d, using each SARS-CoV-2 variant as inoculum. (B) Sequence homology from aa 834 to 863 in Spike among the SARS-CoV2 variants. Asterisks indicate the fully homologous sites among all the variants.

### PA-001 inhibits SARS-CoV-2 propagation and disease progression in vivo by either pretreatment or posttreatment after virus infection

We assessed the antiviral effect of PA-001 in infection animal models. We initially examined the effect of prophylactic treatment with PA-001 before virus inoculation on propagation of infectious SARS-CoV-2 in a Syrian hamster infection model^36,37^ (Fig. 4A-a); Syrian hamsters were pretreated with PA-001 at 0.3, 1.0, or 3.0 mg/kg, or saline as a vehicle control intraperitoneally twice [at 12 and 2 h before virus inoculation (-0.5 and -0.08 day)], followed by intranasal inoculation with SARS-CoV-2 (Delta variant), and then posttreatment with PA-001 at the same concentrations three times [at 12, 24, and 36 h (0.5, 1, and 1.5 day)]. Hamsters were sacrificed at day 2 for collecting lungs to quantify the infectious titer of SARS-CoV-2. SARS-CoV-2 infectious titers detected in these tissues derived from vehicle-administered hamsters were significantly reduced by PA-001 treatment in a dose-dependent manner, by up to 3.5-log (at 3.0 mg/kg of PA-001) (Fig. 4A-b). These data indicate the potential anti-SARS-CoV-2 effect of PA-001 in hamsters by prophylactic treatment.

**Fig. 4.**
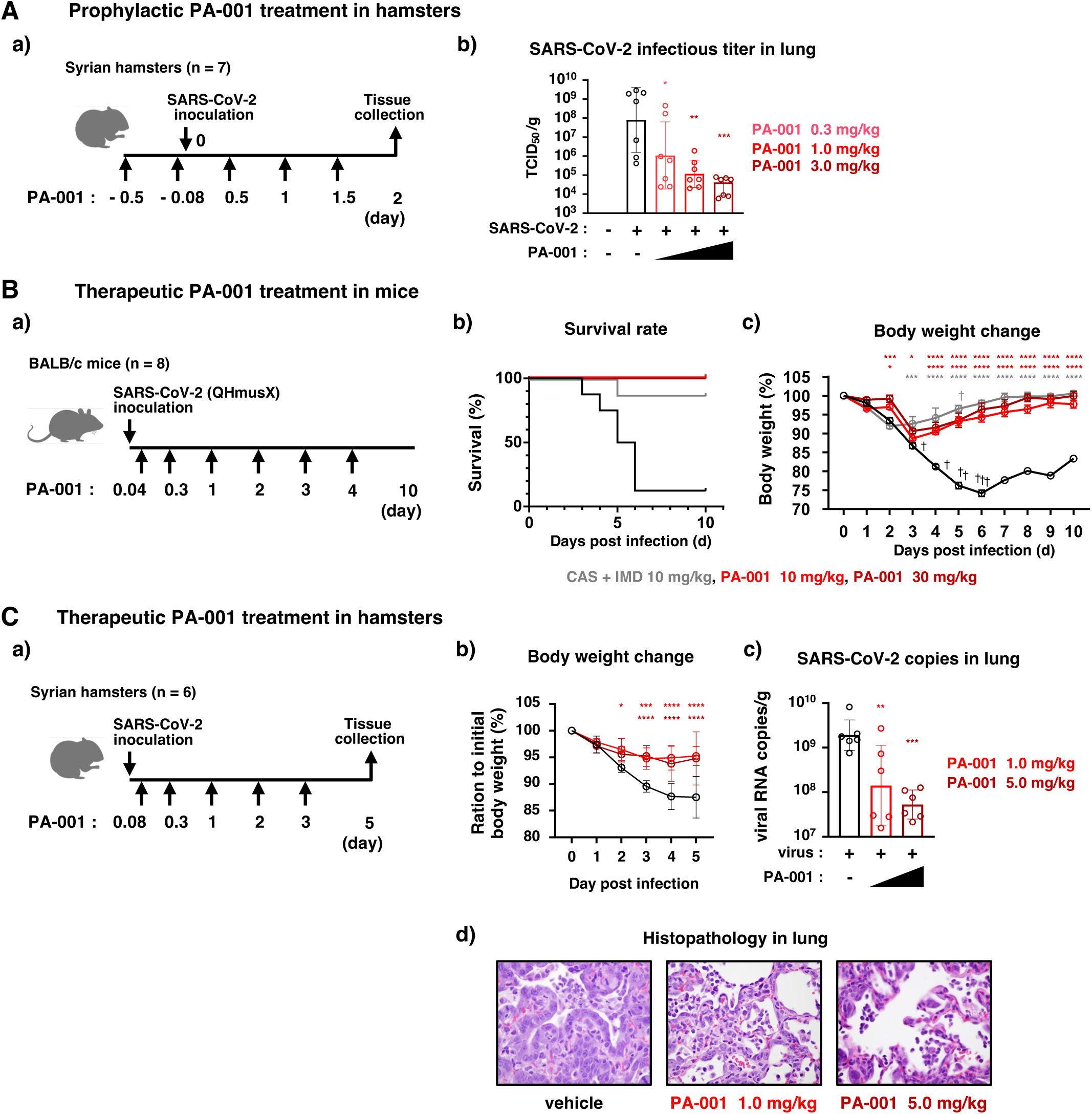
PA-001 inhibited viral propagation and disease progression in mice and hamsters. (A) (a) Scheme for prophylactic treatment with PA-001 to SARS-CoV-2-infected hamsters. Golden Syrian hamsters (n = 7 for infected groups and n = 5 for uninfected group) were intraperitoneally treated with PA-001 (0.3, 1.0, or 3.0 mg/kg) or the vehicle at day -0.5, -0.08, 0.5,1.0, and 1.5 (-12, -2, 12, 24, and 36 h) post-inoculation, and were inoculated with SARS-CoV- 2 (3.48x10^3^ TCID_50_/hamster) intranasally at day 0. Lungs were collected at day 2 post-inoculation. (b) Infectious viral titer in the lungs at day 2 was quantified. Black, vehicle; light red, 0.3 mg/kg PA-001; red, 1.0 mg/kg PA-001; dark red, 3.0 mg/kg PA-001. Ordinary one-way ANOVA followed by Dunnett’s multiple comparisons with the vehicle control was performed (*: p<0.05, **: p<0.01, ***: p<0.001). (B) (a) Scheme for therapeutic treatment with PA-001 to SARS-CoV-2-infected mice. BALB/c mice (n = 8) were intranasally inoculated with mouse-adopted SARS-CoV-2 (QHmusX) (3x10^3^ TCID_50_/mouse) (day 0) and then administered intraperitoneally with PA-001 (10 or 30 mg/kg), CAS + IMD (10 mg/kg), or the vehicle control at day 0.04, 0.3, 1, 2, 3, and 4 (1, 8, 24, 48, 72, and 96 h). The mice were chased to examine the body weight as well as survival ratio. (b, c) Survival rate (b) and body weight (body weight at day 0 as 100%) (c) along with the time course after virus inoculation (day) was plotted. Black, vehicle; gray, 10 mg/kg CAS + IMD; red, 10 mg/kg PA-001; dark red, 30 mg/kg PA-001. Ordinary two-way ANOVA followed by Dunnett’s multiple comparisons with the PBS control was performed (*: p<0.05, **: p<0.01, ***: p<0.001, ****: p<0.0001). (C) (a) Schematic diagram for PA-001 treatment after SARS-CoV-2 infection to hamsters. Golden Syrian hamsters (n = 6) intranasally inoculated with SARS-CoV-2 (2.11x10^5^ TCID_50_/hamster) (day 0) were intraperitoneally treated with PA-001 (1.0 or 5.0 mg/kg) or the vehicle at day 0.08, 0.3, 1.0, 2.0, and 3.0 (2, 8, 24, 48, and 72 h). Lungs were collected at day 5. (b) Body weight was observed daily, defining those at day 0 as 100%. Black, vehicle; red, PA-001 1.0 mg/kg; dark red, PA-001 5.0 mg/kg. Ordinary two-way ANOVA followed by Dunnett’s multiple comparisons with the vehicle control was performed (*: p<0.05, **: p<0.01, ***: p<0.001, ****: p<0.0001). (c) SARS-CoV-2 RNA levels in the lung were quantified at day 5. Ordinary one-way ANOVA followed by Dunnett’s multiple comparisons with the vehicle control was performed (*: p<0.05, **: p<0.01, ***: p<0.001). (d) Histopathology of the lungs at day 5. Lungs from hamsters of each treatment group were stained by HE.

To further evaluate the therapeutic efficacy by posttreatment after infection, we examined the pathogenicity induced by SARS-CoV-2 propagation in mice upon posttreatment with PA-001 after infection; BALB/cCrSlc mice intranasally infected with a mouse-adapted SARS-CoV-2 (QHmusX strain)^38^ were treated intraperitoneally with PA-001 at 10 or 30 mg/kg, saline as a vehicle control, or CAS + IMD as a positive control at 1, 8, 24, 48, 72, and 96 h after virus inoculation (0.04, 0.3, 1, 2, 3, and 4 days), and were chased to measure body weight as well as survival rate daily up to day 10 (Fig. 4B-b, c). Seven mice out of eight treated with the vehicle were dead or reached the humane endpoint (survival rate = 12.5%) at day 6, and body weights of vehicle-treated mice were gradually decreased following SARS-CoV-2 infection to 73-80% of the original until day 5 (Fig. 4B-b, c). In contrast, all mice treated with PA-001 after virus inoculation at either 10 or 30 mg/kg survived (survival rate = 100%), whereas one mouse was dead upon CAS + IMD treatment (survival rate = 87.5%) (Fig. 4B-b). Furthermore, PA-001-treated group showed only a transient body weight reduction by up to around 11 % at day 3, followed by a recovery to the original body weight level at day 10, as the case with CAS + IMD-treated mice (Fig. 4B-c).

We further examined histopathology in the lung as well as body weight and SARS-CoV-2 viral load in the lung in Syrian hamsters posttreated with PA-001 after SARS-CoV-2 infection (Fig. 4C); Hamsters after intranasal SARS-CoV-2 infection were posttreated with PA-001 (1 or 5 mg/kg) or saline at 2, 8, 24, 48, and 72 h and were collected at day 5 to analyze histopathology and viral RNA levels in the lung as well as chased for measuring body weight up to day 5 (Fig. 4C-a). Body weight loss induced by SARS-CoV-2 propagation (to 88% in the vehicle-control group) was significantly recovered by PA-001 posttreatment (to 94% in PA-001-treated group) (Fig. 4C-b). SARS-CoV-2 RNA levels in the lung were reduced by PA-001 treatment in a dose-dependent manner by up to 1.6-log reduction (by 5.0 mg/kg of PA-001) (Fig. 4C-c). Histopathological examination revealed severe pneumonia with consolidation from vehicle-treated hamsters, whereas such severe illness in histopathology was improved by PA-001 treatment, especially at 5 mg/kg (Fig. 4C-d). Thus, PA-001 inhibited SARS-CoV-2 propagation in the lung and suppressed the resultant disease progression by post-treatment after infection in mice and hamsters.

### Combination treatment with PA-001 and remdesivir augments the antiviral effects

Combination treatment with multiple drugs with different mode of action generally enhances the antiviral effect and decreases the risk for selecting drug-resistant variants^39^. We then examined the combination treatment of PA-001 with RDV at various concentrations to evaluate the antiviral effects in cell-based infection assay. As shown in Fig. 5A, RDV reduced SARS-CoV-2 production in a dose-dependent manner by single treatment, and this RDV’s antiviral effects were enhanced by addition of PA-001 (Fig. 5A): e.g., RDV at 1,500 nM and PA-001 at 0.70 nM single treatment reduced viral production to 51% and 43%, respectively, but combination with RDV at 1,500 nM plus PA-001 at 0.70 nM decreased viral production to 11% of the control (Fig. 5A), indicating that PA-001 potentiated the antiviral activity of RDV.

**Fig. 5.**
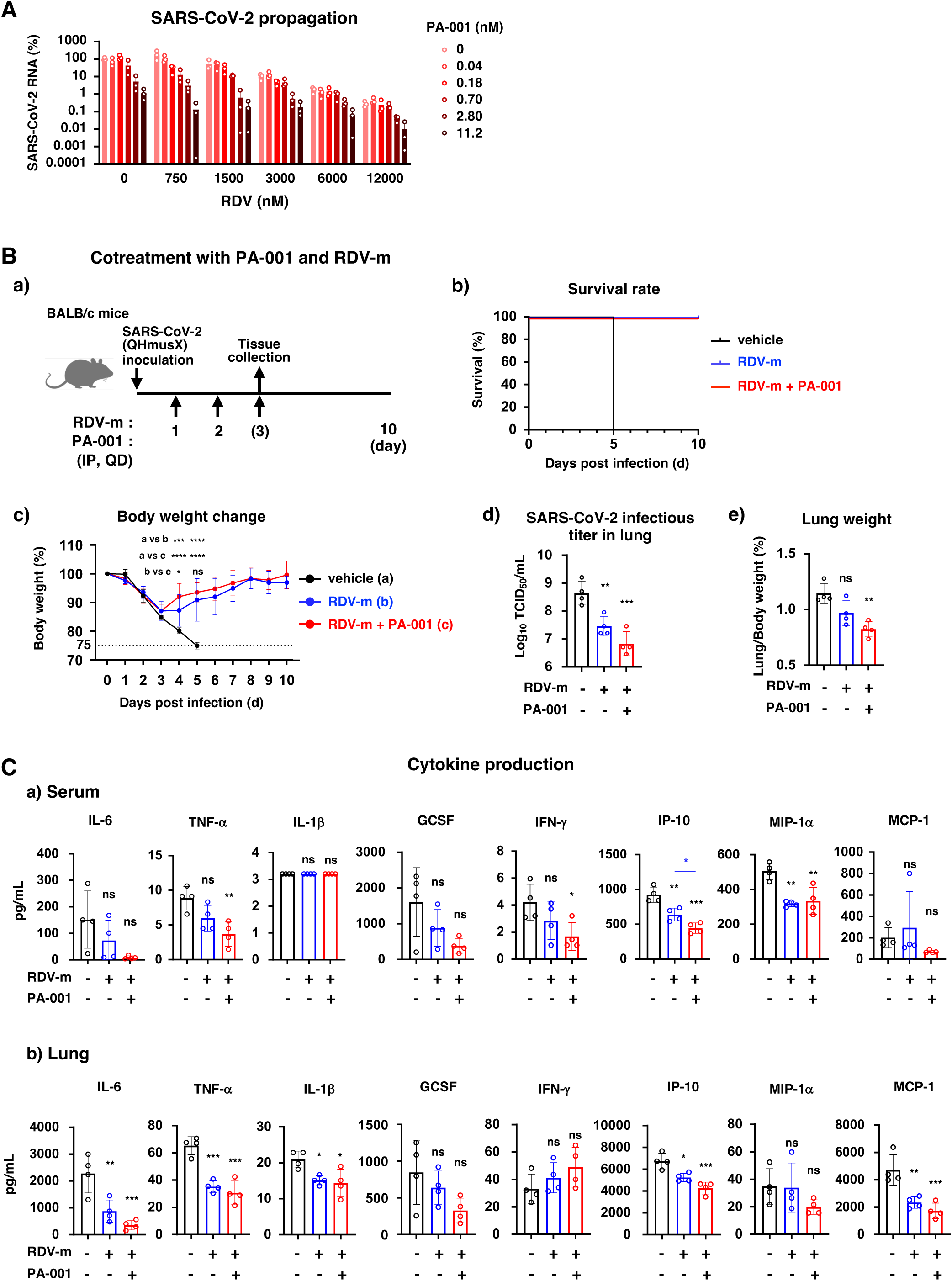
Augmented anti-SARS-CoV-2 profiles by combination treatment with PA-001 and RDV. (A) Combination treatment with PA-001 and RDV in cell-based infection assay. Antiviral activities were evaluated as described in Fig. 1D-d, using SARS-CoV-2-infected VeroE6/TMPRSS2 cells that were treated with combinations of various concentrations of PA-001 (0, 0.04, 0.18, 0.70, 2.8, and 11.2 nM) and RDV (0, 750, 1,500, 3,000, 6,000, and 12,000 nM). (B) (a) Scheme for post-administration of combination with PA-001 and the RDV metabolite, GS-441524 (RDV-m). BALB/c mice were intranasally inoculated with SARS-CoV-2 (2.3x10^3^ TCID_50_/mouse) and then intraperitonially administered with either GS-441524 (25 mg/kg (day 1) and 12.5 mg/kg (days 2, 3)) alone, GS-441524 (25 mg/kg (day 1) and 12.5 mg/kg (days 2, 3)) with PA-001 (26 mg/kg), or the vehicle once daily from day 1 to 3. Mice were euthanized at day 3 without dosing on the day for collecting lungs and sera (n = 4) and day 10 after monitoring survival rate and body weight (n = 5-7). (b, c) Survival rate (b) and body weight (c) until day 10 for each group are shown. Black, vehicle; blue, GS-441524; red, GS-441524 + PA-001. Ordinary two-way ANOVA followed by tukey’s multiple comparisons was performed (*: p<0.05, **: p<0.01, ***: p<0.001, ****: p<0.0001). (d, e) Infectious viral titer in the lung (d) and the lung weight (e, lung weight-to-body weight ratio) at day 3 are shown. Ordinary one-way ANOVA followed by tukey’s multiple comparisons was performed (*: p<0.05, **: p<0.01, ***: p<0.001). (C) Production of the cytokines related with the disease severity, IL-6, TNF-α, IL-1β, GCSF, IFN-γ, IP-10, MIP-1α, and MCP-1, from the serum and the lung at day 3 were analyzed. Ordinary one-way ANOVA followed by tukey’s multiple comparisons was performed (*: p<0.05, **: p<0.01, ***: p<0.001).

We next examined the effect of combination therapy with PA-001 and GS-441524 (RDV-m), the active metabolite of RDV^40^, in SARS-CoV-2 infection mouse model. BALB/cCrSIc mice intranasally inoculated with a mouse-adapted SARS-CoV-2 strain were intraperitoneally treated with RDV-m (25, 12.5, 12.5 mg/kg), RDV-m (25, 12.5, 12.5 mg/kg) + PA-001 (26 mg/kg), or vehicle control (Fig. 5B-a). The vehicle-control group mice showed a continuous decrease in body weight and all the mice died or reached the humane end point at day 5 after infection (Fig. 5B-b, c, black). In contrast, body weight loss of mice treated with RDV-m alone was only transient until day 3 to 4, followed by a gradual recovery, with all the mice survived until day 10 (Fig. 5B-b, c, blue). Consistently, virus infectious titer in the lung was significantly decreased by 1.2-log, although the lung weight, as a marker of tissue damage and inflammation was not significantly improved by single treatment with RDV-m (Fig. 5B-d, e). Combination treatment with RDV-m and PA-001 more rapidly recovered the body weight loss at day 4 to 7 compared with RDV-m single treatment, with no death during the whole experimental period (Fig. 5B-b, c, red), and showed additional reduction in SARS-CoV-2 infectious titer in the lung (by 1.8-log) (Fig. 5B-d, red). Furthermore, lung weight increase by SARS-CoV-2 was significantly improved by RDV-m + PA-001 cotreatment (Fig. 5B-e, red).

We further examined the production profile of cytokines and chemokines, especially those reportedly associated with the COVID-19 severe disease progression, including interleukin (IL)-6, tumor necrosis factor-alpha (TNF-α), IL-1β, granulocyte colony stimulating factor (GCSF), interferon (IFN)-γ, IFNγ-induced protein 10 (IP10), macrophage inflammatory protein 1-alpha (MIP-1α), and monocyte chemoattractant protein-1 (MCP1)^41-44^, in both the serum and the lung of mice at day 3 postinfection. As shown in Fig. 5C, SARS-CoV-2-induced production of IL-6, TNF-α, IL-1β, IP-10, MIP-1α, and MCP-1 at least either in the serum or the lung was significantly inhibited by RDV-m monotreatment, and most of these expressions were more remarkably reduced by combination treatment with RDV-m and PA-001 (Fig. 5C). Furthermore, induction of IP-10 in serum was significantly reduced in PA-001 and RDV-m combination treatment from RDV-m single treatment; Production of TNF-α and IFN-γ in the serum was significantly reduced under only combination treatment with PA-001 and RDV-m, but not single treatment with RDV-m (Fig. 5C). Thus, combination treatment with PA-001 potentiated the anti-SARS-CoV-2 effect of RDV and the resultant disease progression.

### Pharmacokinetics and safety of PA-001 in healthy volunteers

A double-blind, randomized, placebo-controlled phase I study for PA-001 was conducted in 2 parts (Part A and B) to examine the safety, tolerability, and pharmacokinetics of PA-001 in healthy volunteers. Part A was a single-dose sequential-group study with a total of 32 randomized participants. Baseline characteristics were balanced across cohorts, in which mean age of participants was 47 years (range 22–75 years) and the majority of participants were female (59.4%), white (65.6%), and not Hispanic or Latino (62.5%), with a mean BMI of 26.17 kg/m^2^ (Fig. 6A). These volunteers were grouped into five cohorts who were administered intravenously with a single administration of either i) placebo (n = 8), ii) 18 mg PA-001 (n = 6), iii) 64 mg PA-001 (n = 6), or iv, v) 128 mg PA-001 (n = 6 to younger and n = 6 to elderly). Administration with PA-001 at 128 mg had two cohorts, one with younger (median age of 37.2 years, iv) and one with elder volunteers (median age of 69.8 years, v). These cohorts were chased to observe adverse effects as well as to collect the plasma in a time course up to 48 h post-administration for measuring the concentration of PA- 001. As a result, PA-001 was well-tolerated, showing no severe adverse effects related with treatment (treatment emergent adverse effects, TEAEs) in all individuals (Fig. 6B and Supplementary Table 4A). Observed events were mild, including headache (1 subject in Placebo, 2 subjects in 64 mg PA-001) and vomiting (1 subject in 64 mg PA-001) (Fig. 6B and Supplementary Table 4A). Pharmacokinetics profiles were presented with plasma concentrations against the time course up to 48 h after single dosing (Fig. 6C), and pharmacokinetic parameters of PA-001 at each dose calculated based on these profiles (Fig. 6D and Supplementary Table 5). Dose proportional increases in the peak concentrations and exposures of PA-001 were observed across the evaluated dose range (Fig. 6C). PA-001 was readily detectable in plasma at 0.5 h after administration with an average maximal concentrations (C_max_) of up to 18,300 ng/mL (9,610 nM) at 128 mg treatment, and was gradually declined, with 3.61-4.79 h as half-life (T_1/2_). Given the IC_50_ of 0.93-2.2 nM in the cell-based infection assay against the Omicron strains, BA.1.18 and XBB.1.16 (Supplementary Table 3), 128 mg PA-001 group exhibited more than 4,300-10,300-fold concentration of IC_50_ in plasma at peak (C_max_) at 1 h after administration (Fig. 6C). The overall exposure of PA-001 (AUC_0-tlast_) was up to 74,200 ng·h /mL at 128 mg administration (younger).

**Fig. 6.**
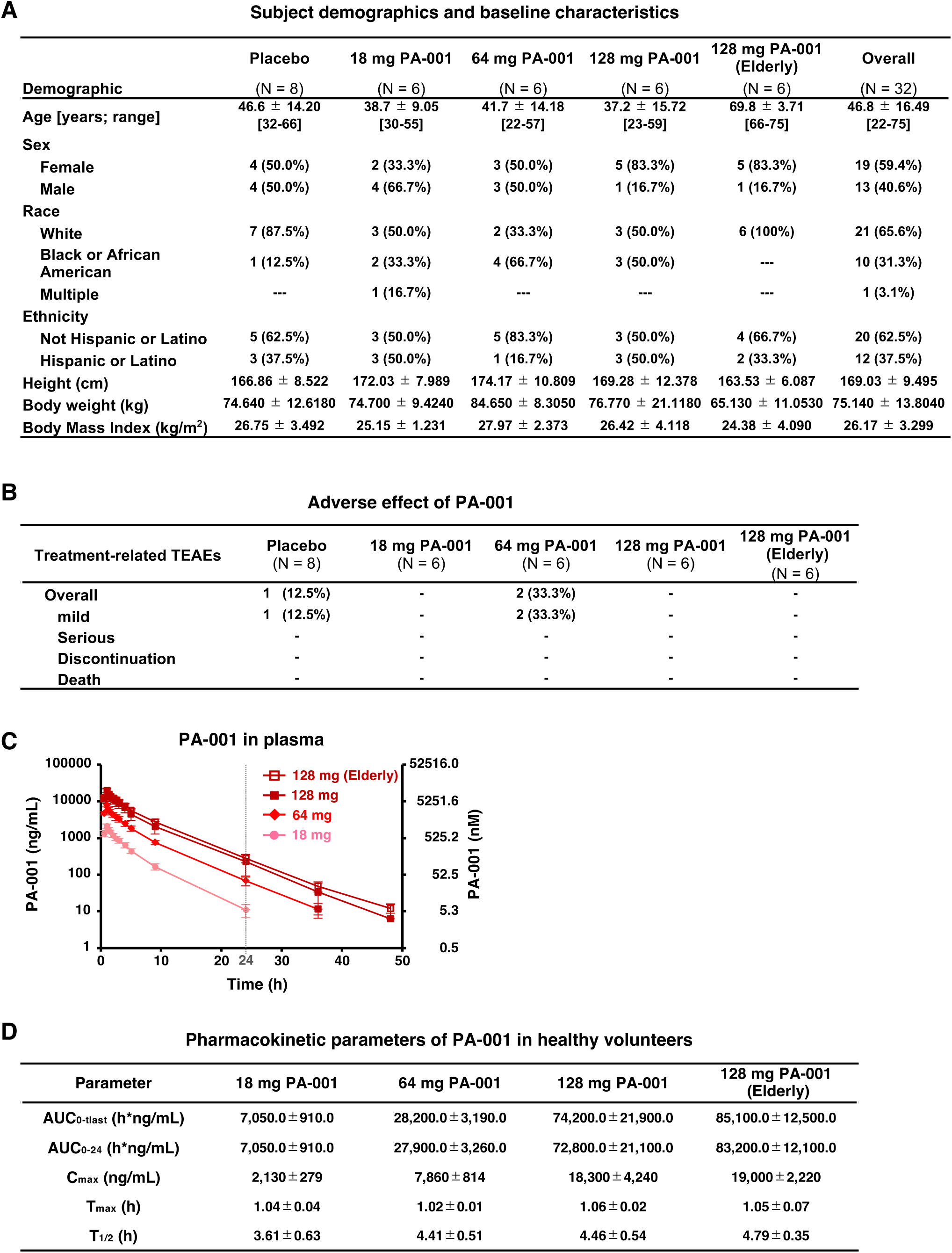
Safety and pharmacokinetics profiles of PA-001 single administration to healthy volunteers. (A) Baseline characteristics of 32 participants in the phase I trial for PA-001. The indicated profiles are presented as mean ± standard deviation for each cohort group (Placebo, 18 mg PA-001, 64 mg PA-001, 128 mg PA-001, and 128 mg PA-001 to elderly adults). Numbers of participants for each group are also shown (n = 6 or 8 for each group). (B) Number of participants as well as percentage showing the treatment-related treatment emergent adverse effects (TEAEs) as indicated for each cohort group. (C) Plasma concentrations of PA-001 for each cohort (closed circle, 18 mg; closed diamond, 64 mg; closed square, 128 mg; open square,128 mg to elderly adults) in the single-dose sequential-group study (Part A) are plotted against the time post intravenous infusions in healthy subjects. (D) Pharmacokinetic parameters calculated based on the profile shown in (C). AUC_0-tlast_, area under the concentration-time curve from time 0 to the endpoint; AUC_0-24_, area under the concentration-time curve from 0 to 24 h; C_max_, maximum concentration; T_max_, time for maximum concentration; T_1/2_, half-life. Data are represented as mean ± standard deviation.

Part B was a multi-dose sequential-group study, in which total 15 participants were randomized into three groups who received multiple dosing of either placebo (n = 4), 64 mg PA-001 (n = 5), or 128 mg PA-001 (n = 6) once a day until day 5 (Supplementary Table 6). The mean age of participants was 37 years (range 24–60 years); Eight (53.3%) males and seven (46.7%) females, with a mean BMI of 27.29 kg/m^2^. The majority (66.7%) of subjects were White (Supplementary Table 6). Pharmacokinetics parameters on day 1 and day 5 (Supplementary Table 7) were similar to those of single dosing in part A (Fig. 6D and Supplementary Table 5). Following multiple doses of 64 and 128 mg for 5 days, AUC_0-τ_ and C_max_ increased in a supra proportional manner with respect to dose. The AR_AUC_ (Accumulation ratio based on AUC_0-_τ) was 1.01 for the 64 mg dose level and 1.03 for the 128 mg dose level (Supplementary Table 7B). Because there was no accumulation, steady state concentrations of PA-001 were achieved from the onset of the multiple dosing period. No serious adverse effects related with multi-dosing of PA-001 was observed, although seven (46.7%) subjects had at least 1 mild TEAE, of whom 2 received 64 mg PA-001 (Grade 1), 4 received 128 mg PA-001 (3 of Grade 1, 1 of Grade 2), and 1 received placebo (Grade 1) (Supplementary Table 4B). Three (20.0%) subjects had Grade 1 TEAE possibly related to study treatment (2 received 64 mg, 1 received 128 mg), but all these TEAEs had resolved by the end of the study (Supplementary Table 4B). There were no deaths, serious adverse effects, or discontinuations because of adverse effects.

Thus, PA-001 provided the sustained plasma concentration in human individuals above those required for antiviral activity, with preferable safety profiles.

## Discussion

In this study, we identified a 13-amino acid macrocyclic peptide, PA-001, specifically binding to FPPR on the S2 subunit of the SARS-CoV-2 Spike protein, which has not been targeted by antibodies or antiviral agents so far. Despite extensive efforts of cryo-EM analysis, we were unable to obtain a high-resolution structure of the PA-001/Spike complex, probably due to the characteristic of FPPR, having high conformational flexibility, as mentioned below^7,45,46^. Alternatively, to demonstrate the binding site, we created SARS-CoV-2 variants more than 100-fold resistance to PA-001 and found an amino acid substitution cluster within residues 836-862, which resides at FPPR, upstream of FP. Mutations in this region completely abolished Spike’s capacity to bind to PA-001, confirming this segment as the PA-001-binding site. In the pre-fusion form, FPPR is positioned on the surface of Spike trimer between the convex CTD1 and CTD2, forming a shallow depression at the surface. FPPR is suggested to control the RBD movement between its up and down conformations^4^: FPPR is structured and clamps RBD in the down conformation of the pre-fusion form through interaction of Asp614 (in RBD) with Lys854 (in FPPR), whereas it moves out and unstructured in the RBD down conformation^4,47-49^. Upon ACE2-RBD engagement and the following S1 release, FPPR, together with the adjacent FP, undergoes an overall translocation during the conformational rearrangement to be positioned at the top of the S2 trimer and form a cone-shaped structure, which functions to be inserted into the target cellular membrane. The binding of PA-001 to FPPR at the pre-fusion form (Fig. 2A and F) would sterically interfere with this mobility of FPPR toward the formation of the cone-shaped structure (intermediation form; Supplementary Fig. 1) that is required for the fusion process. These findings highlight FPPR as a previously unexplored but potential target for antiviral drug development.

Previously developed Spike-targeting therapeutics largely act through antibodies directed against S1 such as casirivimab and imdevimab, amubarvimab and romlusevimab, bamlanivimab, bamlanivimab and etesevimab, sotrovimab, tixagevimab and cilgavimab, and bebtelovimab^14,15^. These antibodies primarily exert anti-SARS-CoV-2 activity by neutralization of infection through blocking Spike-ACE2 interaction and/or by engaging Fc-mediated effector functions that promote viral clearance. However, their neutralization activity has been severely impaired or completely lost by accumulated mutations in RBD of the S1 subunit among the recent Omicron sublineages. The NTD in S1, another target of antibodies, also has high sequence variability. In contrast, the S2 region is more conserved in sequence and represents a target for developing broadly active antivirals. S2-targeting antibodies reported so far mainly recognize the region downstream of the S2′ site (residues 811-825, previously referred to as FP, but recently considered an intermediate loop) or the stem helices upstream of HR2, however, they had relatively modest antiviral potency, likely due to the limited surface accessibility by antibodies as well as low immunogenicity^17,50,51^. To overcome such limitations, non-antibody modalities have been applied so far, such as peptides and aptamers, which have smaller molecular size that can bind to regions inaccessible to conventional antibodies. HR2-mimetic peptides, such as EK1 and HY3000, designed to interfere with the HR1-HR2 interaction during the conformational change to the post-fusion form, have shown broad antiviral activity across SARS-CoV-2 variants, with reported original IC_50_ at 500-3,000 nM, and their activities were further optimized through the lipid conjugation^17,18^. S2-targeting DNA aptamers have also been reported, though most of their antiviral activities in infection models remains to be characterized^52^. Among all these S2-directed agents, PA-001 exhibited one of the most potent anti-SARS-CoV-2 activities reported to date (IC_50_ = 0.23–2.9 nM of PA-001 v.s. 2468 nM of EK1^18^ and 580 nM of HY3000^20^. The second key characteristics of PA-001 lies in its target region, FPPR, as a highly conserved region with minimal variations. The third notable feature of PA-001 is its 13-residue macrocyclic structure, markedly smaller than the previously reported mimicking peptides mentioned above (typically more than 30 residues), making it accessible to FPPR positioned in shallow depression at the surface of Spike pre-fusion form. In addition, its cyclic architecture generally confers improved proteolytic stability, reduced immunogenicity, and favorable pharmacokinetics.

So far, macrocyclic peptides have been applied to target SARS-CoV-2 Spike to develop anti-SARS-CoV-2 agents, majority of which target RBD in S1^53-57^. Among them, in vivo anti-SARS-CoV-2 effect was shown for three series of peptides, 6L3-1F3P11hR (against RBD), multivalent bicyclic peptides (against RBD), and S-20-1 (targeting both RBD and HR1)^54,56,57^. However, antiviral activity of S-20-1 was relatively mild, with 540-10,230 nM as IC_50_ in cell-based infection assay, and that of 6L3-1F3P11hR varied at 1-332 nM depending on the target SARS-CoV-2 variants, consistent with the property of RBD-targeting agents. None of them have been reported to be administered in human. In addition to its distinct target (FPPR), PA-001 shows an advantage among these macrocyclic peptides with its high activity in vivo and administration profile in human.

In infection mouse and hamster models, PA-001 effectively inhibited SARS-CoV-2-induced disease progression when administered before or after infection, underscoring its potential for both prophylactic and therapeutic use. Additionally, combination treatment with PA-001 with RDV elevated therapeutic profiles, highlighting its well compatibility with already-approved antivirals. This is consistent with that PA-001 has a distinct mode of action from those of any clinically approved anti-SARS-CoV-2 agents so far. Given the demand for effective anti-COVID-19 therapies, especially for immunocompromised patients who respond poorly to current treatments and evolve drug-resistance^1-3^, peptide-based antivirals with different mode of action would provide an additional choice for COVID-19 therapy. Encouragingly, phase I clinical trial of PA-001 in 32 healthy volunteers demonstrated well tolerability (up to 128 mg administration of PA-001) without serious adverse events, regardless of age. In the single-dose cohort, the C_max_ reached 18,300 ng/mL [9,610 nM, more than 4,300-fold higher than IC_50_ (0.93–2.2 nM) against Omicron variants] with half-life of 4.46 h, and drug exposure was sufficient to maintain concentrations above the cellular IC_50_ for up to 24 h post-dose in Groups of 128 mg administration. Collectively, these findings suggest that PA-001 achieves robust and sustained systemic exposure, supporting a favorable pharmacokinetic profile. Therefore, our data raise PA-001 as a promising anti-SARS-CoV-2 drug that targets S2 and inhibits fusion process, and forthcoming clinical trials will assess its efficacy in SARS-CoV-2-infected patients.

In summary, our study shows the usefulness of macrocyclic peptides as a potential therapeutic modality against COVID-19, that demonstrates FPPR in the S2 region as a druggable target to inhibit virus fusion process. The strong antiviral potency warrants PA-001 as a candidate seed for the development of an anti-SARS-CoV-2 fusion inhibitor.

## Methods

### Synthesis of PA-001

PA-001 was synthesized using standard Fmoc solid-phase peptide synthesis (SPPS), as described in the Supplementary Notes. After coupling of all amino acids, the N-terminus was deprotected and subsequently chloroacetylated on resin, followed by global deprotection using a standard deprotection cocktail containing trifluoroacetic acid (TFA). The linear peptide was precipitated by the adding the deprotection solution to more than a 10-fold excess of diethyl ether and hexane. The resulting crude peptide pellet was resuspended in diethyl ether and re-pelleted three times. After the final wash, the pellet was dried and dissolved in DMSO/H₂O, followed by the addition of triethylamine to induce intramolecular cyclization via thioether bond formation between the cysteine thiol and the N-terminal chloroacetyl group. Upon completion of cyclization, the cyclic peptide was purified by standard reverse-phase HPLC. The molecular mass was confirmed by single-quadrupole LC/MS (LCMS-2020 system, Shimadzu).

### Cell culture

VeroE6/TMPRSS2 cells, a VeroE6 cell line overexpressing transmembrane protease, serine 2 (TMPRSS2) and highly susceptible to SARS-CoV-2 infection^25^, were cultured in Dulbecco’s modified Eagle’s medium (DMEM; Life Technologies) supplemented with 10% fetal bovine serum (FBS; Cell Culture Bioscience), 10 units/mL penicillin, 10 mg/mL streptomycin, 10 mM HEPES (pH 7.4), and 1 mg/mL G418 (Nacalai) at 37°C in 5% CO_2_. During the infection assay, 10% FBS was replaced with 2% FBS, with removing G418. HEK293T cells were cultured in DMEM supplemented with 10% FBS (heat-inactivated) and 50 μg/mL gentamicin at 37°C in 5% CO_2_.

### Reagents

Anti-SARS-CoV-2 Spike S1 antibodies, casirivimab (CAS) and imdevimab (IMD), were cloned into human IgG1 and kappa expression vectors and co-transfected into Expi293F cells (Thermo Fisher Scientific) as described previously^58^. Mutant spike proteins were generated by introducing substitutions into expression vectors coding codon-optimized spike trimeric protein^59^ by PCR, and expressed with Expi293F cells. Antibodies and mutant spike proteins were purified from the culture supernatant using Protein G columns (Thermo Fisher Scientific) and TALON columns (Clontech), respectively, and dialyzed against PBS. Nucleotide analogs, RDV and its active metabolite, GS-441524 (RDV-m), were purchased from Chemscene Inc (CS-0028115) and Cayman Chemical (30469), respectively. PA-001 and HA tagged PA-001 were produced by PeptiDream Inc. DMSO was purchased from Sigma-Aldrich. Mefloquine was purchased from Tokyo Chemical Industry.

### Cell-based infection assay

SARS-CoV-2 was handled in a biosafety level 3 (BSL3). We used the SARS-CoV-2 Wk-521 (Wuhan type), QK002 (Alpha), TY8-612 (Beta), TY7-503 (Gamma), TY11-927 (Delta), TY38-873 (Omicron; BA.1.18), and TY41-984 (Omicron; XBB.1.16) strains, clinical isolates from a COVID-19 patient^25^. Virus infectious titers were measured by inoculating cells with a 10-fold serial dilution of virus and cytopathology measured to calculate TCID_50_/mL^25^. For the infection assay, VeroE6/TMPRSS2 cells were inoculated with virus at an MOI of 0.003 for 1 h and washed to remove unbound virus. Cells were cultured for an additional 24, 36, and 48 h prior to detection of extracellular viral RNA or intracellular viral N protein (24 or 36 h), and cytopathic effects (CPE) (48 h), respectively. Compounds were added during virus inoculation (1 h) and incubation after washing (24, 36, or 48 h), or incubation after washing only.

### Isolation and quantification of viral RNA

Viral RNA was extracted with RNeasy mini kit (QIAGEN) from cells and MagMax Viral/Pathogen II Nucleic Acid Isolation kit (Thermo Fisher Scientific) or QIAamp Viral RNA mini kit (QIAGEN) from culture supernatant. Viral RNA was quantified by real time RT-PCR analysis with a one-step qRT-PCR kit (THUNDERBIRD Probe One-step qRT-PCR kit, TOYOBO) using 5’-ACAGGTACGTTAATAGTTAATAGCGT-3’, 5’- ATATTGCAGCAGTACGCACACA-3’ as a primer set, and 5’-FAM-ACACTAGCCATCCTTACTGCGCTTCG-TAMRA-3’ (E-set) as a probe^60^.

### Detection of viral proteins

Viral proteins were detected using rabbit anti-SARS-CoV N antibody^61^ and mouse anti-SARS-CoV-2 spike antibody (abcam, ab273433) as primary antibodies, with anti-rabbit IgG-HRP (Cell Signaling, 7074) and anti-mouse IgG-HRP (Cell Signaling, 7076) as secondary antibodies by immunoblot analyses as previously reported^62^.

### Virus-cell binding assay

SARS-CoV-2 preincubated with the indicated compounds for 30 min was exposed to VeroE6/TMPRSS2 cells at an MOI of 0.01 at 4°C for 15 min. After extensive wash, total RNA was extracted from cells and viral RNA was quantified by real time RT-PCR to measure cell-bound virus, as described ^63^.

### PA-001-Spike binding assay

Binding of HA-tagged PA-001 to SARS-CoV-2 Spike proteins was quantified by ELISA as follows: Recombinant wild type (from Sino biological, 40589-V08B1, 40591-V08B1, and 40590-V08B) or mutated Spike proteins were immobilized on the plates, and then incubated with PA-001-HA at various concentrations together with or without excess amount of non-tagged PA-001, used as a binding competitor. After washing with PBST, horseradish peroxidase-labeled mouse anti-HA tag antibody (MBL Life Science, M180-7, 1:5000 or 1:2500) was added, and visualized with reacting with a substrate solution (SUMITOMO BAKELITE) to detect absorbance at 450 nm with a microplate reader (TECAN).

### Cell membrane fusion assay

The S-ACE2 interaction-mediated cell-cell fusion assay was performed essentially based on the previous report^29^, with some modifications. Spike proteins expressed in HEK293T cells mediate the fusion with VeroE6/TMPRSS2 cells, and the cell-cell fusion was quantified based on the measurement of luminescence intensity derived from reconstituted NanoLuc reporter (composed of HiBiT stably expressed in VeroE6/TMPRSS2 cells and LgBiT transiently expressed in HEK293T cells). HEK293T cells were co-transfected with the plasmid expressing Spike in combination with the HaloTag-inserted pBiT1.1-N[TK/LgBiT] plasmid (Promega) using Lipofectamine 3000 Transfection Reagent according to the manufacturer’s instruction (Thermo Fisher Scientific). At 48 h post-transfection, the cells were co-incubated with VeroE6/TMPRSS2/HiBiT cells in the presence or absence of varying concentrations of PA-001 for 1 h. The media were replaced with Opti-MEM containing 0.5% FBS and the substrate of NanoLuc from Nano-Glo Live Cell Assay System (Promega) to measure luminescence.

### Generation of PA-001 resistant viruses

To obtain PA-001-resistant SARS-CoV-2 variants, SARS-CoV-2 (WK-521) was inoculated to VeroE6/TMPRSS2 cells at an MOI of 0.003, followed by incubation of the cells in the presence of 1,000 nM PA-001 for 48 h and recovery of culture supernatant. The 1:1 diluted culture supernatant was then inoculated to VeroE6/TMPRSS2 cells to repeat the same procedure until the second passage. The 1:10 diluted culture supernatant was inoculated to the cells in the presence of 10 μM PA-001, followed by recovery of culture supernatant at 24 h later, which was repeated for the three additional passages. Virus was further amplified with two additional passages in the absence of PA-001.

### SARS-CoV-2 pseudovirus infection assay

SARS-CoV-2 pseudoviruses were produced using vesicular stomatitis virus (VSV)-based pseudovirus system, using the expression plasmid for the G-deficient VSV carrying the luciferase gene and that encoding SARS-CoV-2 Spike derived from the Wuhan type Wk-521 or the Omicron sublineages, JN.1 or XFG, essentially as previously described^64,65^. VeroE6/TMPRSS2 cells were inoculated with the SARS-CoV-2 pseudovirus in the presence or absence of various concentrations of the indicated compounds, peptide, or antibodies followed by incubation with or without the compounds, peptide, or antibodies for 24 h to measure intracellular luciferase activity with Luciferase Assay System (Promega), as described^62^.

### Sequence analysis of PA-001 resistant viruses

cDNA libraries were generated from RNA extracted from SARS-CoV-2 isolates according to the ARTIC nCoV-2019 sequencing protocol v3 (LoCost) (https://www.protocols.io/view/ncov-2019-sequencing-protocol-v3-locost-bp2l6n26rgqe/v3). Whole-genome sequencing was performed using the MinION platform (Oxford Nanopore Technologies), followed by consensus sequence generation using the Medaka pipeline (https://github.com/nanoporetech/medaka). Lineage and clade assignments, as well as mutation identification, were conducted using the Pangolin COVID-19 Lineage Assigner (https://pangolin.cog-uk.io/) and Nextclade (https://clades.nextstrain.org/).

### Pharmacokinetic analysis of PA-001 in mice

The study was conducted under accreditation by the Association for Assessment and Accreditation of Laboratory Animal Care (AAALAC) with approval of Institutional Animal Care and Use Committee (IACUC, protocol No. PK-T-07182021). Fasted female BALB/c mice (8 weeks old, Si Bei Fu Laboratory Animal Technology) were intraperitoneally administered with PA-001 at the dose of 10 mg/kg, and were sacrificed to collect blood at 1, 7, and 24 h post-administration. The blood was centrifuged to prepare plasma, mixed with acetonitrile containing formic acid, and centrifuged. The supernatant was injected onto LC/MS/MS system (LCMS-8060, Shimadzu) to determine plasma concentrations of PA-001.

### SARS-CoV-2 infection in vivo

For infection to the mouse model in Fig. 4B and Fig. 5B and C, all the experiments were approved by the Animal Care and Use Committee of the National Institute of Infectious Diseases in Japan, and handled in BSL3 animal facilities (ABSL3) that were certified by the Japan Health Sciences Foundation according to the guidelines of this committee. BALB/c mice (13 weeks old, female, Japan SLC) were maintained for 3 weeks in the specific pathogen-free facility and then transferred to the ABSL3. After a week of acclimation to the facility, the mice (17 weeks old) were anesthetized by intraperitoneal injection with a mixture of 0.1 ml/10 g body weight of 1.0 mg ketamine and 0.02 mg xylazine. Animals were then inoculated intranasally with 3 x 10^3^ TCID_50_ (30 µl, 5 LD_50_/mouse, 100% lethal dose) QHmusX, a mouse-adopted SARS-CoV-2 clone^38^. Body weight was measured daily for 10 days (n = 10–11 per group), and animals were sacrificed at 3 and 10 days post-inoculation to analyze viral replication, cytokine expression, and disease pathology (n = 4 or 6–7 per group). Clinical signs were observed up until 10 days post-inoculation. The humane endpoint was defined as the appearance of clinically diagnostic signs of respiratory stress, including respiratory distress and more than 25% weight loss. Animals were euthanized under anesthesia with an overdose of isoflurane if severe disease symptoms or weight loss were observed. In Fig. 4B, PA-001 was administered intraperitoneally at a dose of 10 or 30 mg/kg, at 0.04, 0.3, 1, 2, 3, and 5 days post-inoculation. The CAS + IMD-treatment group received the cocktail CAS + IMD antobodies (10 mg/kg) at 1 h post-inoculation. The vehicle control group was administered with PBS. In, Fig. 5B and C, RDV-m was administered intraperitoneally at a dose of 25 mg/kg (on day 1 post-infection) and 12.5 mg/kg (on days 2 and 3 post-infection), with a volume of 6.25 mL/kg per dose, once daily. Concurrently, PA-001 was administered intraperitoneally at a dose of 26 mg/kg, with a volume of 10 mL/kg per dose, once daily. The control group was administered with PBS.

Infection studies to the hamster model shown in Fig. 4A and C were conducted in the animal BSL-3 laboratory, accredited by AAALAC, under IACUC-approved protocol (22-032, 16103.01). In Fig. 4A, seven-week-old male Syrian hamsters (Envigo) were intranasally inoculated with 3.48 x 10^3^ TCID_50_ of SARS-CoV-2 (Delta variant, BEI Resources, NR-56116) under anesthesia with ketamine/xylazine at day 0. PA-001 was intraperitoneally administered to hamsters twice daily at 0.3, 1, or 3 mg/kg, or PBS as a vehicle control, at 12 h and 2 h prior to infection and 12 h, 24 h, and 36 h following infection (-0.5, -0.08, 0.5, 1, and 1.5 day post infection). Lung in the hamsters were collected at 2 days post-infection to examine the infectious titer of SARS-CoV-2. In Fig. 4C, Syrian Golden hamsters (female, 8 week-old, Charles Reiver Laboratories) were intranasally infected with 2.11 x 10^5^ TCID_50_ SARS-COV2 (USA_WA1/2020, CDC, kindly provided by UTMB Galveston), and then administered intraperitonially with either PBS as a vehicle control or PA-001 1 or 5 mg/kg (n=6/group) (7.5 mL/kg) at 2, 8, 24, 48 and 72 h post-infection. Body weight and clinical observation were evaluated every day, and the lungs were collected at five day postinfection. The left lung lobes were used for quantifying SARS-CoV-2 RNA by real time RT-PCR, whereas the right lung lobes were fixed with 10% neutral buffered formalin for histopathological evaluation.

### Viral titration from mouse and hamster tissues

Lung tissue samples from BALB/c mice were collected at the time of postmortem examination and stored at -80°C. Tissue homogenates (10% w/v) were prepared in Dulbecco’s Modified Eagle medium (DMEM) containing 2% FBS, 50 IU/ml penicillin G, and 50 μg/ml streptomycin, and samples were inoculated onto VeroE6/TMPRSS2 cell cultures, which were then examined for observing cytopathic effects (CPEs) for 5 days to determine the viral infectivity titers (TCID_50_) as described above.

Lung tissue samples from Syrian hamsters were homogenized and centrifuged to collect the supernatant. VeroE6/TMPRSS2 cells were treated with the supernatant to evaluate CPEs for 4 days to determine the viral infectivity titers with the Read-Muench formula^66^.

### Detection of inflammatory cytokines and chemokines

Homogenized mouse lung tissue samples (10% w/v) were diluted 1:1 in cell extraction buffer [10 mM Tris (pH 7.4), 100 mM NaCl, 1 mM EDTA, 1 mM EGTA, 1 mM NaF, 20 mM Na_4_P_2_O_7_, 2 mM Na_3_VO_4_, 1% Triton X-100, 10% glycerol, 0.1% SDS, and 0.5% deoxycholate (BioSource International, Camarillo, CA)], incubated for 10 min on ice with vortexing, irradiated for 10 min with UV-C light to inactivate infectious virus. Cytokine and chemokine levels were measured with mouse cytokine–chemokine magnetic bead panel 96-well plate assay kit (Milliplex MAP kit, Merck Millipore), detected on a Luminex 200 instrument with xPONENT software (Merck Millipore), according to the manufacturer’s instruction.

### Phase I study

#### 1. Study design and oversight

This phase I, double-blind, randomized, placebo-controlled, single- and multiple-ascending dose study of the safety, tolerability, and pharmacokinetics in healthy and elderly subjects was conducted at a single clinical site, Fortrea Clinical Research Unit Inc. The protocol (PA-001-101) was approved by Salus institutional review board (IRB). Study began in Sep 2024 and was completed in Jun 2025 (from FSFV to LPLV). All participants provided written informed consent before enrollment.

#### 2. Study Design

This study was conducted in 2 parts of Part A and B, as shown below.

Part A comprised a single-dose, sequential-group study of PA-001. A total of up to 40 subjects were planned to be studied in up to 4 groups (Groups A1-4, PA-001 doses were 18, 64, 128 and 128 mg, respectively, with each group comprising 8 subjects. In each group, 6 subjects were to receive a single dose of PA-001 and 2 subjects were to receive a single dose of placebo. Age range was 18 to 65 years for Group A1-3, and >65 years for Group A4. Doses of PA-001 or placebo were administered in accordance with a randomization schedule on the morning of Day 1, after a fast of at least 10 h. Sentinel dosing: Groups A1 to A4 were divided into 2 subgroups, with each subgroup being dosed at least 24 h apart. The first subgroup comprised 2 subjects, with 1 subject receiving PA-001 and 1 subject receiving placebo. The second subgroup comprised 6 subjects, with 5 subjects receiving PA-001 and 1 subject receiving placebo. Continuation to dose the remaining 6 subjects was at the investigator’s discretion.

Part B comprised a multiple-dose, sequential group study. Part B was permitted to start after completion of Group A3 (in Part A), at a dose level less than or equal to that administered in Groups A1 to A3. A total of 16 subjects were planned to be studied in 2 groups, with each group comprising 8 subjects. In each group, 6 subjects were to receive PA-001, and 2 subjects were to receive placebo once per day until day 5. PA-001 or placebo were administered in accordance with a randomization schedule after a fast of at least 10 h.

**I**nclusion criteria and exclusion criteria were described in Supplementary Notes.

#### 3. Safety assessments

The condition of each subject was monitored from the time of signing the ICF to final discharge from the study. Subjects were observed for any signs or symptoms and asked about their condition by open questioning, such as “How have you been feeling since you were last asked?” at least once each day while resident at the study site and at each study visit. Subjects were also encouraged to spontaneously report adverse effects occurring at any other time during the study. All nonserious adverse effects, whether reported by the subject voluntarily or on questioning, or noted on physical examination, were recorded from initiation of study treatment until study completion. Serious adverse effects were recorded from the time the subject signs the ICF until study completion. The nature, time of onset, duration, and severity were documented, together with an investigator’s (or designee’s) opinion of the relationship to study treatment. Adverse events recorded during the study will be followed up, where possible, until resolution or until the unresolved adverse effects were judged by the investigator (or designee) to have stabilized. This was completed at the investigator’s (or designee’s) discretion.

#### 4. Administration

In Part A, subjects received a single 50mL intravenous infusion over 1 h (± 2 min) containing 18 mg (Group A1), 64 mg (Group A2), or 128 mg (Groups A3 and A4) of PA-001 or placebo on day 1. In part B, subjects received multiple 50mL intravenous infusions over 1 h (± 2 min) containing 64 mg (Group B1) or 128 mg (Group B2) of PA-001 or placeboonce per day on day 1 through 5, inclusive.

#### 5. Analysis of PA-001 concentrations

Concentration of PA-001 in plasma and urine samples was measured with liquid chromatography tandem mass spectrometry (LC/MS/MS, QTRAP® 5500 System, SCIEX).

## Statistics and reproducibility

Bar graphs are presented as mean ± standard deviation with individual data sets, and experiments were repeated in at least triplicates or more unless indicated otherwise. In this study, GraphPad Prism (v.10.5.0) was used for the ordinary one-way ANOVA followed by Dunnett’s multiple comparisons with control (Fig. 2C-a, Fig. 4A-b, and Fig. 4C-c) or followed by tukey’s multiple comparisons (Fig. 5B-d, e, and C-a, b), and the ordinary two-way ANOVA followed by Dunnett’s multiple comparisons with control (Fig. 4B-c, Fig. 4C-b) or followed by tukey’s multiple comparisons (Fig. 2F-b and Fig. 5B-c).

## Acknowledgments

We appreciate Drs. Setsuo Hasegawa (Pharma Spur), Akihito Shimoi (SNBL), Takuma Yonemura, Takeshi Chiyoda, Asuka Yamamoto (Sumida Hospital) for valuable scientific advice based on clinical research experience, which contributed to the rationale for the starting dose in the Phase I study. We also thank Dr. Keiichi Masuya for PA-001 generation. We are grateful to Drs. Dai Izawa, Midori Ozaki, Takiko Yoshida, for their technical assistance. We thank the members of the Management Department of Biosafety and Laboratory Animals for support with the BSL3 facility. We also appreciate leadership of Dr. Gene Voskuhl (Fortrea Clinical Research) to operate Phase I study. This study was supported by Japan Agency for Medical Research and Development (AMED) under grand numbers JP25fk0310525 (K.W.), JP23fk0108590 (K.W.), JP25ak0101271 (K.W.), 25wm0325079 (K.W.), JP253fa727002 (N.Nagata, T.S.), JP25wm0125008 (T.S.), JP25fk0108732 (T.S.), JP223fa627003 (T.S.), JP22fk0108521 (M.M.); JSPS/MEXT under KAKENHI grant numbers JP24K18457 (H.Ohashi), JP24K02290 (K.W.), JP25K21782 (K.W.), JP25K21767 (K.W.), JP25K02691 (K.W.); JST MIRAI program grant number JPMJMI22G1 (K.W.); Takeda Science Foundation (K.W.).

## Author Contributions

Experimental systems were developed by M.O., T.S., N.Nagata and K.W. Compound screening was supervised by M.O. and P.C.R. M.O., K.N.,T.N., K.S.,N.Nakamura and J.Y. performed screening for lead compound of PA-001. M.O., K.M. and H.Kurasaki optimized peptides and identified PA-001. Molecular biological and cell-based experiments were performed by H.Ohashi, Y.H., K.Shionoya, T.K., H.Ogawa, and K.W. Animal experiments using mice were designed and performed by N.IY., N.Nagata, and H.Kitamura, and those using hamsters were designed and managed by T.K., K.N. and H.Kitamura. Peptide synthesis was performed by M.O., H.Kurasaki, K.N., T.N., J.Y., K.Sudo, N.Nakamura, and K.M. Non-clinical ADME analyses was conducted by K.Y., Y.S. and T.H. Preparing materials was managed by H.Kurasaki, S.M., T.M., T.H., and S.I. Experimental design was developed by N.Nagata, M.O., P.C.R., T.S., and K.W. Phase I clinical trial was designed and managed by S.I., N.K., and M.M. Original draft was written by H.Ohashi, N.Nagata, S.M., H.Kitamura, and K.W. with critical discussion by T.K, M.O., H.Kurasaki, N.IY., Y.H., S.M., K.Shionoya, K.N., T.N., J.Y., K.Sudo., N.Nakamura, K.M., H.Ogawa, K.Yoshida, Y.S., T.M., T.H., S.I., Y.T., N.K., P.C.R., M.M., and T.S. Fundings were acquired by H.Ohashi, M.M., T.S., N.Nagata, and K.W. The project was supervised by H.Kitamura and K.W.

## Data Availability

Source data are provided with this paper.

