## Supplementary Notes, Supplementary Table 1-7, Supplementary Figure 1-3 for "A macrocyclic peptide-based fusion inhibitor targeting SARS-CoV-2 Spike S2 subunit"

**Contents**

**Supplementary Notes**
**Supplementary Table 1-7**
**Supplementary Figure 1-3**

#### Supplementary Notes

##### Solid-phase peptide synthesis, cyclization, and purification of PA-001

HMPB-MBHA resin (Merck, 0.85 mmol/g) was loaded with Fmoc-D-Glu(OtBu)-OH using DIC and DMAP (4 equiv, 4 equiv, and 0.1–0.5 equiv per equivalent of resin, respectively) in DMF/DCM (1:40, v/v) at room temperature for 60 min. Using the resulting Fmoc-D-Glu(OtBu)-HMPB-MBHA resin, the target peptide was synthesized according to a general solid-phase peptide synthesis protocol. Peptide assembly was carried out on a Liberty Blue automated peptide synthesizer (CEM) following the manufacturer's standard procedures. For the incorporation of each amino acid residue, Fmoc-AA, DIC, and Oxyma Pure (4.2 equiv, 8 equiv, and 4 equiv, respectively, relative to the resin) were employed, and the coupling reaction was performed once in DMF at 75°C for 10 min. However, modified coupling conditions were applied for specific residues: the first residue (Fmoc-MeE(tBu)-OH) and the second residue (Fmoc-MeNle-OH) were coupled twice at 75 °C for 30 min, and the twelfth residue (Fmoc-Cys(Trt)-OH) was coupled once at 50 °C for 15 min. Fmoc deprotection was generally carried out by treatment with a 20% piperidine solution in DMF at 75 °C for 3 min. However, for the first through sixth residues, the deprotection reaction was performed twice at 25 °C for 5 min and 10 min. Introduction of the chloroacetyl group was achieved by first removing the Fmoc group from the α-amino group of the resin-bound peptide obtained in the previous step using the method described above. Subsequently, a 0.1 M solution of chloroacetic acid in DMF (5 equiv), a 0.1 M solution of HATU in DMF (5 equiv), and a 0.2 M solution of DIEA in DMF (10 equiv) were added to the resin. The mixture was then shaken at room temperature for 30 min. Side-chain deprotection and cleavage of the peptide from the solid-phase resin were performed by washing the resin obtained after the chloroacetylation step with DMF and methylene chloride, followed by drying under reduced pressure. The resin was then transferred to a reaction vessel, and cocktail-A (TFA/$H_2O$/TIS (Triisopropylsilane)/DODT (3,6-Dioxa-1,8-octanedithiol), 92.5:2.5:2.5:2.5, v/v/v/v) was added. The mixture was shaken at 25 °C for 10 min. The reaction solution was collected by filtration through a frit. The solid-phase resin remaining in the reaction vessel was subsequently shaken again with the cleavage cocktail, and the resulting solution was collected through the frit and combined with the initial filtrate. Addition of the combined filtrate to an excess of diethyl ether/hexane (1:1, v/v) precooled to 0 °C resulted in the formation of a cloudy precipitate. The mixture was centrifuged at 9,000 rpm for 2 min, and the supernatant was decanted. The resulting solid was washed with a small amount of diethyl ether precooled to 0 °C and then dried under reduced pressure. The obtained solid was used directly in the subsequent cyclization reaction. Peptide cyclization was performed by dissolving the peptide in DMSO/$H_2O$ (9:1, v/v) to give a final peptide concentration of 5 mM, calculated based on the initial loading of the solid-phase resin. Triethylamine (10 equiv) was added, and the reaction mixture was stirred at 25 °C for 17 h. Upon completion, the reaction solution was concentrated under reduced pressure using an EZ-2 Elite evaporator. The crude product was purified by preparative reverse-phase HPLC under the following conditions: column, Waters Xbridge C18 (5 μm, 50 × 250 mm); mobile phase, A = 0.1% TFA in $H_2O$ and B = 0.1% TFA in MeCN; column temperature, 50 °C; gradient (%B), 14.4–11.1% over 0.1 min, 11.1% over 4.9 min, 11.1–14.4% over 2 min, 14.4–38.8% over 3 min, 38.8–43.9% over 15 min, and 43.9–60% over the subsequent 3 min; flow rate, 120 mL/min. After lyophilizing the solution, the product was purified again by preparative reverse phase HPLC under the following conditions: column, Kinetex EVO C18 (30 × 150 mm); mobile phase, A = 1.0% AcOH in $H_2O$ and B = 1.0% AcOH in MeCN; column temperature, 40 °C; gradient (%B), 8–33% over 3min, 33–38% over 8min, 38–60% over 1 min; flow rate, 45 mL/min. The purity of the target product was determined from the peak area ratio of the LC/MS chromatogram monitored at 225 nm and was found to be 99.4%. Analytical LC/MS was performed under the following conditions: retention time, 5.40 min; column, Kinetex EVO C18 (2.6 μm, 2.1 × 150 mm, 100 Å); mobile phase, A = 0.025% TFA in $H_2O$ and B = 0.025% TFA in MeCN; column temperature, 60 °C; gradient (%B), 20–60% over 7.15 min, 60–95% over 0.30 min, followed by 95% for 1.55 min; flow rate, 0.5 mL/min. ESI-MS (positive mode) gave an observed m/z value of 922.92 corresponding to $[M + 2H]^{2+}$.

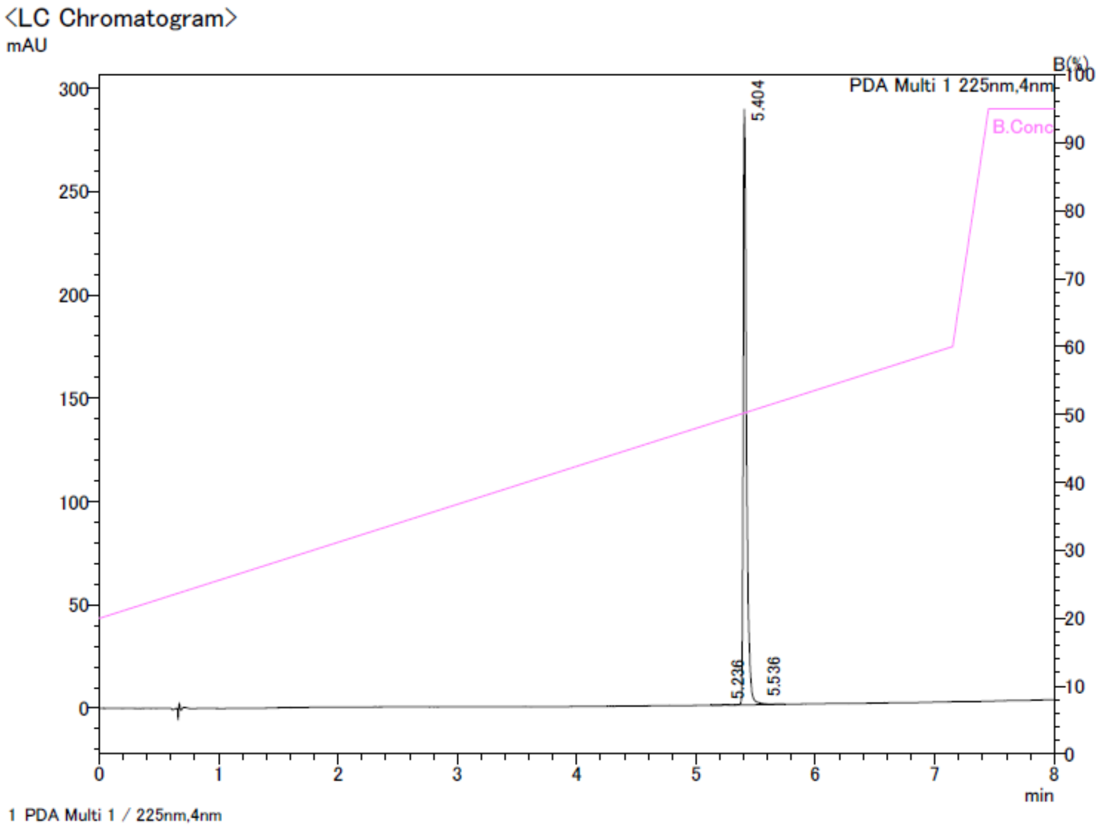


PDA Ch1 225nm

| peak# | tR | Area | Area(%) | Height | Height(%) |
|---|---|---|---|---|---|
| 1 | 5.236 | 1049 | 0.193 | 255 | 0.088 |
| 2 | 5.404 | 540191 | 99.378 | 288474 | 99.654 |
| 3 | 5.536 | 2331 | 0.429 | 747 | 0.258 |
| Total | | 543571 | 100.000 | 289475 | 100.000 |

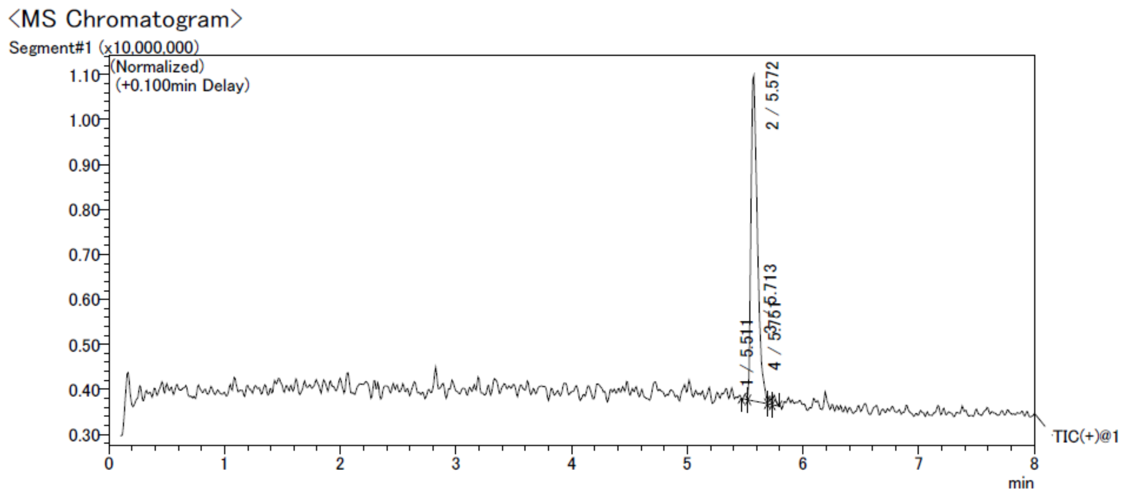


MS Peak Table TIC

| peak# | tR | Area | Area(%) | Height | Height(%) | Peak m/z | Base Peak Intensity |
|---|---|---|---|---|---|---|---|
| 1 | 5.511 | 262871 | 0.918 | 137170 | 1.767 | 1424.86 | 25872 |
| 2 | 5.572 | 27722815 | 96.782 | 7227745 | 93.118 | 923.02 | 5559083 |
| 3 | 5.713 | 304923 | 1.065 | 173104 | 2.230 | 922.92 | 117564 |
| 4 | 5.751 | 353862 | 1.235 | 223868 | 2.884 | 923.05 | 71219 |
| Total | | 28644472 | 100.000 | 7761887 | 100.000 | | |

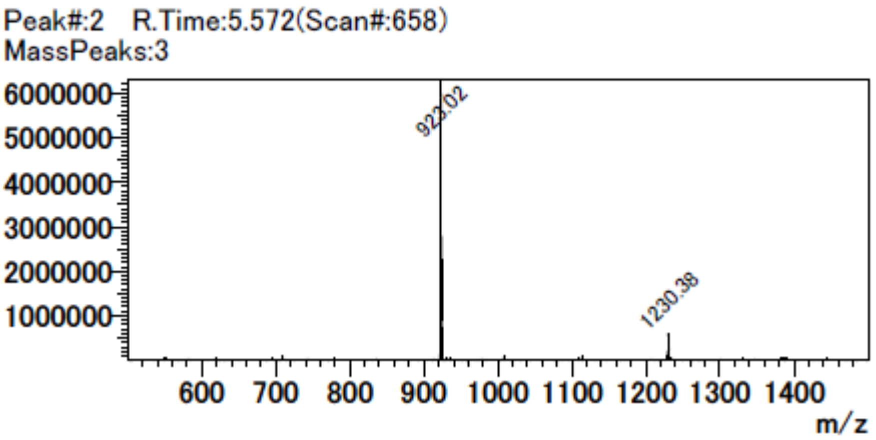


**Eligibility criteria of clinical trial**

**Inclusion criteria**

Subjects must satisfy all of the following criteria at screening unless otherwise stated:

1. Groups A1 to A3, B1, and B2 only: Males or females, of any race, between 18 and 65 years of age, inclusive.
2. Group A4 (and optionally A5) only: Males or females, of any race, > 65 years of age.
3. Body mass index between 18.0 and 32.0 kg/m$^2$, inclusive.
4. In good health, or have stable, chronic, non life threatening medical conditions, determined by no clinically significant findings from medical history, 12 lead ECG, vital signs measurements, and clinical laboratory evaluations (congenital nonhemolytic hyperbilirubinemia [eg, suspicion of Gilbert's syndrome based on total and direct bilirubin] is not acceptable) at screening and/or check in and from the physical examination at check in, as assessed by the investigator (or

designee).

5. Females will not be pregnant or lactating, and females of childbearing potential and males will agree to use contraception.
6. Able to comprehend and willing to sign an ICF and to abide by the study restrictions.

**Exclusion criteria**

Subjects will be excluded from the study if they satisfy any of the following criteria at screening unless otherwise stated:

Medical Conditions

1. Significant history or clinical manifestation of any metabolic, allergic, dermatological, hepatic, renal, hematological, pulmonary, cardiovascular, gastrointestinal, neurological, respiratory, endocrine, or psychiatric disorder that, in the opinion of the investigator (or designee), could impact subject safety or the objectives of the study.
   Note: Subjects who have stable, chronic, non life threatening medical conditions, including, but not limited to, hypertension, hypo- or hyper-thyroidism, or osteoarthritis, or postmenopausal state, for at least 3 months are permitted to be enrolled, providing that, in the opinion of the investigator (or designee), it will not impact subject safety or the objectives of the study.
2. Have signs and symptoms of any other liver disease, except nonalcoholic fatty liver disease, or any of the following, as determined from clinical laboratory evaluations:
   a. alanine aminotransferase or aspartate aminotransferase >1.25 × the upper limit of normal (ULN)
   b. alkaline phosphatase or total bilirubin >1.5 × ULN, except if known or suspected Gilbert's syndrome
   c. lactate dehydrogenase > 1.5 × ULN
   d. glomerular filtration rate <60 mL/min, based on Chronic Kidney Disease Epidemiology Collaboration 2021 formula.
   e. platelet count <150,000/$mm^3$
   f. hemoglobin <12 g/dL for males or <11 g/dL for females.
3. History of significant hypersensitivity, intolerance, or allergy to any drug compound, food, or other substance, as determined by the investigator (or designee).
4. Positive hepatitis panel or positive human immunodeficiency virus test.
5. Positive SARS-CoV-2 test at screening or check in.
6. Any of the following:
   a. QT interval corrected for heart rate (QTc) using Fridericia's method (QTcF) > 450 ms in males or > 470 ms in females confirmed by calculating the mean of the original value and 2 repeats
   b. QRS duration > 110 ms confirmed by calculating the mean of the original value and 2 repeats
   c. PR interval ≥ 220 ms confirmed by calculating the mean of the original value and 2 repeats
   d. findings which would make QTc measurements difficult or QTc data uninterpretable
   e. history of additional risk factors for torsades de pointes (eg, heart failure, clinically significant

hypokalemia, family history of long QT syndrome).

Prior/Concomitant Therapy

7. Administration of a COVID-19 vaccine in the past 30 days prior to dosing.
8. Use or intend to use any of the following that could potentially impact subject safety or the objectives of the study, as determined by the investigator (or designee):
   a. any prescription medications/products other than hormone replacement therapy; oral, implantable, transdermal, injectable, or intrauterine contraceptives; or medications/products for the treatment of hypertension, hypo- or hyper-thyroidism, osteoarthritis, or those used for postmenopausal state within 14 days prior to dosing
   b. any nonprescription medications/products including vitamins, minerals, and phyto-therapeutic/herbal/plant derived preparations within 7 days prior to dosing
   c. any medications that are substrates of cytochrome P450 (CYP) 2A6, organic anion transporter (OAT) 1, OATPB1, OATP1B3, organic cation transporter 2 (OCT2), multidrug and toxin extrusion protein 2-K (MATE2-K), or bile salt export pump (BSEP).4

Prior/Concurrent Clinical Study Experience

9. Participation in a clinical study involving administration of an investigational drug (new chemical entity) in the past 30 days or 5 half lives of that drug prior to dosing, whichever is longer.
10. Have previously completed or withdrawn from this study or any other study investigating PA-001 and have previously received PA-001.

Diet and Lifestyle

11. Alcohol consumption of > 21 units per week for males and > 14 units for females. One unit of alcohol equals 12 oz (360 mL) beer, 1½ oz (45 mL) liquor, or 5 oz (150 mL) wine.
12. Positive urine drug screen at screening, or positive alcohol test result or positive urine drug screen at check in.
13. History of alcoholism or drug/chemical abuse within 2 years prior to check in, as defined by the current Diagnostic and Statistical Manual of Mental Disorders.
14. Use of tobacco or nicotine containing products within 3 months prior to check in, or positive cotinine at screening or check in.
15. Ingestion of Seville orange, or grapefruit containing foods or beverages within 7 days prior to check in. Ingestion of poppy seed containing foods or beverages within 3 days prior to check in.

Other Exclusions

16. Receipt of blood products within 2 months prior to check in.
17. Donation of blood from 3 months prior to screening, plasma from 2 weeks prior to screening, or platelets from 6 weeks prior to screening.
18. Poor peripheral venous access.
19. Subjects who, in the opinion of the investigator (or designee), should not participate in this study.

#### Supplementary Tables

**Supplementary Table 1.**
**Binding kinetics of PA-001 with recombinant SARS-CoV-2 Spike**

| Parameter | Unit | Value |
|---|---|---|
| $k_a$ | $Ms^{-1}$ | $4.37 \times 10^{5}$ |
| $k_d$ | $s^{-1}$ | $7.86 \times 10^{-4}$ |
| $K_D$ | M | $1.80 \times 10^{-9}$ |

$k_a$: Association rate constant.
$k_d$: Dissociation rate constant.
$K_D$: Equilibrium dissociation constant.

**Supplementary Table 2.**

**Pharmacokinetic parameters of PA-001 in BALB/c mice**

| Parameter | Unit | Value |
|---|---|---|
| $t_{1/2}$ | h | 1.7 |
| $T_{max}$ | h | 1.0 |
| $C_{max}$ | ng/mL | 76,667 |
| $AUC_{0-24}$ | ng×h/mL | 282,236 |

$t_{1/2}$: Elimination half-life from plasma

$T_{max}$: Time to reach $C_{max}$

$C_{max}$: Maximum plasma concentration

$AUC_{0-24}$: Area under the curve of a plasma concentration (ng/mL) versus time (h), indicating the total exposure amount of PA-001 until 24 h post administration

**Supplementary Table 3.**

**Antiviral activities of PA-001, CAS/IMD, and RDV to SARS-CoV-2 variants**

| SARS-CoV-2 variants | $IC_{50}$ PA-001 (nM) | $IC_{50}$ CAS/IMD (ng/mL) | $IC_{50}$ RDV (nM) | $IC_{90}$ PA-001 (nM) | $IC_{90}$ CAS/IMD (ng/mL) | $IC_{90}$ RDV (nM) |
|---|---|---|---|---|---|---|
| Wuhan | 0.58 | 8.2 | 2,000 | 1.7 | 27 | 4,400 |
| Alpha | 0.23 | 1.1 | 350 | 1.3 | 6.6 | 2,000 |
| Beta | 0.71 | 2.1 | 1,900 | 1.9 | 7.3 | 3,900 |
| Gamma | 0.28 | 15 | 430 | 1.3 | 32 | 2,000 |
| Delta | 2.9 | 2.5 | 1,400 | 9.3 | 12 | 5,000 |
| Omicron BA.1.18 | 0.93 | > 1,000 | 710 | 2.4 | > 1,000 | 3,100 |
| Omicron XBB.1.16 | 2.2 | > 1,000 | 570 | 9.4 | > 1,000 | 1,600 |

$IC_{50}$: 50% inhibitory concentration

$IC_{90}$: 90% inhibitory concentration

**Supplementary Table 4.**

**Treatment emergent adverse events (TEAEs) in the PA-001 single dosing (Part A) (A) and multiple dosing (Part B) (B) to healthy volunteers**

**A**

| | **Placebo** (N = 8) | **18 mg PA-001** (N = 6) | **64 mg PA-001** (N = 6) | **128 mg PA-001** (N = 6) | **128 mg PA-001 (Elderly)** (N = 6) | **Overall** (N = 32) |
|---|---|---|---|---|---|---|
| TEAEs | | | | | | |
| Overall | 1 (12.5%) | 1 (16.7%) | 3 (50.0%) | — — — | — — — | 5 (15.6%) |
| Serious | — — — | — — — | — — — | — — — | — — — | — — — |
| Leading to Discontinuation | — — — | — — — | — — — | — — — | — — — | — — — |
| Leading to Death | — — — | — — — | — — — | — — — | — — — | — — — |
| Severity | | | | | | |
| Grade 1 | 1 (12.5%) | 1 (16.7%) | 3 (50.0%) | — — — | — — — | 5 (15.6%) |
| Grade 2 | — — — | — — — | 1 (16.7%) | — — — | — — — | 1 (3.1%) |
| Grade 3 | — — — | — — — | — — — | — — — | — — — | — — — |
| Grade 4 | — — — | — — — | — — — | — — — | — — — | — — — |
| Grade 5 | — — — | — — — | — — — | — — — | — — — | — — — |
| Treatment-related TEAEs | | | | | | |
| Overall | 1 (12.5%) | — — — | 2 (33.3%) | — — — | — — — | 3 (9.4%) |
| Serious | — — — | — — — | — — — | — — — | — — — | — — — |
| Leading to Discontinuation | — — — | — — — | — — — | — — — | — — — | — — — |
| Leading to Death | — — — | — — — | — — — | — — — | — — — | — — — |
| Severity | | | | | | |
| Grade 1 | 1 (12.5%) | — — — | 2 (33.3%) | — — — | — — — | 3 (9.4%) |
| Grade 2 | — — — | — — — | — — — | — — — | — — — | — — — |
| Grade 3 | — — — | — — — | — — — | — — — | — — — | — — — |
| Grade 4 | — — — | — — — | — — — | — — — | — — — | — — — |
| Grade 5 | — — — | — — — | — — — | — — — | — — — | — — — |

**B**

| | **Placebo** (N = 4) | **64 mg PA-001** (N = 5) | **128 mg PA-001** (N = 6) | **Overall** (N = 15) |
|---|---|---|---|---|
| TEAEs | | | | |
| Overall | 1 (25.0%) | 2 (40.0%) | 4 (66.7%) | 7 (46.7%) |
| Serious | — — — | — — — | — — — | — — — |
| Leading to Discontinuation | — — — | — — — | — — — | — — — |
| Leading to Death | — — — | — — — | — — — | — — — |
| Severity | | | | |
| Grade 1 | 1 (25.0%) | 2 (40.0%) | 3 (50.0%) | 6 (40.0%) |
| Grade 2 | — — — | — — — | 1 (16.7%) | 1 (6.7%) |
| Grade 3 | — — — | — — — | — — — | — — — |
| Grade 4 | — — — | — — — | — — — | — — — |
| Grade 5 | — — — | — — — | — — — | — — — |
| Treatment-related TEAEs | | | | |
| Overall | — — — | 2 (40.0%) | 1 (16.7%) | 3 (20.0%) |
| Serious | — — — | — — — | — — — | — — — |
| Leading to Discontinuation | — — — | — — — | — — — | — — — |
| Leading to Death | — — — | — — — | — — — | — — — |

| | | | | |
|---|---|---|---|---|
| Severity | | | | |
| Grade 1 | − − − | 2 (40.0%) | 1 (16.7%) | 3 (20.0%) |
| Grade 2 | − − − | − − − | − − − | − − − |
| Grade 3 | − − − | − − − | − − − | − − − |
| Grade 4 | − − − | − − − | − − − | − − − |
| Grade 5 | − − − | − − − | − − − | − − − |

The nS (%) statistics presented

nS: number of subjects with an adverse event

%: percentage of subjects with an adverse event (nS/N x 100)

N: number of subjects

Severity grades: 1 = mild, 2 = moderate, 3 = severe, 4 = life-threatening, 5 = fetal

Adverse events were assigned severity grade using the Division of Allergy and Infectious Diseases (DAIDS) Table for Grading the Severity of Adult and Pediatric Adverse Events, version 2.1.

A treatment-emergent adverse event (TEAE) was defined as an adverse event that started during or after the start of the first infusion, or started prior to the start of the first infusion and increased in severity after the start of the first infusion.

A treatment-related TEAE was defined as a TEAE with a relationship of possibly related or related to the study treatment, as determined by the investigator.

Where a subject experienced multiple TEAEs with the same preferred term for the same treatment, this was counted as 1 TEAE for that treatment under the maximum severity recorded.

**Supplementary Table 5.**

**Pharmacokinetic parameters for the single dosing of PA-001 to healthy volunteers (Part A)**

| Matrix | Parameter | 18 mg PA-001 (N = 6) | 64 mg PA-001 (N = 6) | 128 mg PA-001 (N = 6) | 128 mg PA-001 (Elderly) (N = 6) |
|---|---|---|---|---|---|
| Plasma | $AUC_{0\text{-}tlast}$ (h•ng/mL) | 7,050 ± 910 | 28,200 ± 3,190 | 74,200 ± 21,900 | 85,100 ± 12,500 |
| | $AUC_{0\text{-}24}$ (h•ng/mL) | 7,050 ± 910 | 27,900 ± 3,260 | 72,800 ± 21,100 | 83,200 ± 12,100 |
| | $AUC_{0\text{-}\infty}$ (h•ng/mL) | 7,110 ± 920 | 28,300 ± 3,160 | 74,300 ± 21,900 | 85,200 ± 12,500 |
| | $\%AUC_{extrap}$ (%) | 0.831 ± 0.371 | 0.278 ± 0.174 | 0.163 ± 0.0422 | 0.0992 ± 0.0309 |
| | $C_{max}$ (ng/mL) | 2,130 ± 279 | 7,860 ± 814 | 18,300 ± 4,240 | 19,000 ± 2,220 |
| | $t_{max}$ (h) | 1.04 ± 0.0375 | 1.02 ± 0.00678 | 1.06 ± 0.0172 | 1.05 ± 0.0653 |
| | $t_{1/2}$ (h) | 3.61 ± 0.362 | 4.41 ± 0.514 | 4.46 ± 0.542 | 4.79 ± 0.347 |
| | CL (L/h) | 2.57 ± 0.372 | 2.28 ± 0.241 | 1.88 ± 0.657 | 1.53 ± 0.220 |
| | $V_{ss}$ (L) | 10.1 ± 1.56 | 10.7 ± 1.95 | 8.98 ± 2.75 | 8.38 ± 1.06 |
| | $CL_R$ (L/h) | 0.985 ± 0.557 | 1.24 ± 0.336 | 1.43 ± 0.493 | 0.718 ± 0.203 |
| | $DAUC_{0\text{-}tlast}$ (h•ng/mL/mg) | 392 ± 50.6 | 441 ± 49.8 | 580 ± 171 | 665 ± 97.3 |
| | $DAUC_{0\text{-}24}$ (h•ng/mL/mg) | 392 ± 50.6 | 435 ± 51.0 | 569 ± 165 | 650 ± 94.4 |
| | $DAUC_{0\text{-}\infty}$ (h•ng/mL/mg) | 395 ± 51.1 | 442 ± 49.4 | 581 ± 171 | 666 ± 97.4 |
| | $DC_{max}$ (ng/mL/mg) | 118 ± 15.5 | 123 ± 12.7 | 143 ± 33.1 | 149 ± 17.4 |
| Urine | $Ae_{0\text{-}48h}$ (mg) | 7.08 ± 4.27 | 34.9 ± 9.58 | 97.8 ± 12.8 | 61.2 ± 20.0 |
| | $fe_{0\text{-}48h}$ (%) | 39.4 ± 23.7 | 54.5 ± 15.0 | 76.4 ± 9.97 | 47.8 ± 15.6 |

mean ± SD statistics presented

$AUC_{0\text{-}tlast}$: area under the curve from time 0 to the time of the last quantifiable concentration

$AUC_{0\text{-}24}$: area under the curve over the time interval 0 to 24 h post dose

$AUC_{0\text{-}\infty}$: area under the curve from time 0 extrapolated to infinity

$\%AUC_{extrap}$: percentage of area under the curve due to extrapolation from the last quantifiable concentration to infinity

$C_{max}$: maximum observed concentration

$t_{max}$: time of the maximum observed concentration

$t_{1/2}$: apparent terminal elimination half life

CL: total clearance

$V_{ss}$: volume of distribution of steady state

$CL_R$: renal clearance

$DAUC_{0\text{-}tlast}$: $AUC_{0\text{-}tlast}$ normalized by dose administered

$DAUC_{0\text{-}24}$: $AUC_{0\text{-}24}$ normalized by dose administered

$DAUC_{0\text{-}\infty}$: $AUC_{0\text{-}\infty}$ normalized by dose administered

$DC_{max}$: $C_{max}$ normalized by dose administered

$Ae_{0\text{-}48h}$: amount of the dose administered recovered over the time interval 0 to 48 h

$fe_{0\text{-}48h}$: percentage of the dose administered recovered over the time interval 0 to 48 h

**Supplementary Table 6.**

**Screening demographic data of healthy volunteers for multiple dosing of PA-001 (Part B)**

| Demographic | **Placebo** (N = 4) | **64 mg PA-001** (N = 5) | **128 mg PA-001** (N = 6) | **Overall** (N = 15) |
|---|---|---|---|---|
| Age (years) | 30.5 ± 6.61 [24 − 38] | 40.6 ± 14.48 [26 − 60] | 38.3 ± 9.07 [31 − 53] | 37.0 ± 10.78 [24 − 60] |
| Sex | | | | |
| Male | 2 (50.0%) | 4 (80.0%) | 2 (33.3%) | 8 (53.3%) |
| Female | 2 (50.0%) | 1 (20.0%) | 4 (66.7%) | 7 (46.7%) |
| Race | | | | |
| White | 3 (75.0%) | 4 (80.0%) | 3 (50.0%) | 10 (66.7%) |
| Black or African American | 1 (25.0%) | 1 (20.0%) | 3 (50.0%) | 5 (33.3%) |
| Ethnicity | | | | |
| Not Hispanic or Latino | 2 (50.0%) | 2 (40.0%) | 4 (66.7%) | 8 (53.3%) |
| Hispanic or Latino | 2 (50.0%) | 3 (60.0%) | 2 (33.3%) | 7 (46.7%) |
| Height (cm) | 167.30 ± 10.908 | 173.20 ± 1.977 | 168.08 ± 6.306 | 169.58 ± 6.924 |
| Body Weight (kg) | 68.750 ± 11.095 | 85.020 ± 8.049 | 79.700 ± 7.610 | 78.550 ± 10.418 |
| Body Mass Index (kg/$m^2$) | 24.58 ± 3.551 | 28.34 ± 2.615 | 28.22 ± 2.501 | 27.29 ± 3.124 |

n = number of subjects with valid observation

N = number of subjects

SD = standard deviation

% = percentage of subjects with valid observations (n/N x 100)

For continuous data, mean ± SD statistics presented

For categorical data, n (%) statistics presented

Body mass index (kg/$m^2$) = body weight (kg) / height (m)

**Supplementary Table 7.**

**Pharmacokinetic parameters on day 1 (A) and 5 (B) for multiple dosing of PA-001 (Part B)**

**A**

| Parameter | **64 mg PA-001** (N = 5) | **128 mg PA-001** (N = 6) |
|---|---|---|
| $AUC_{0\text{-tlast}}$ (h•ng/mL) | $26{,}400 \pm 4{,}490$ | $59{,}200 \pm 11{,}100$ |
| $AUC_{0\text{-}\tau}$ (h•ng/mL) | $26{,}400 \pm 4{,}500$ | $59{,}300 \pm 11{,}100$ |
| $AUC_{0\text{-}\infty}$ (h•ng/mL) | $26{,}900 \pm 4{,}800$ | $60{,}200 \pm 11{,}500$ |
| $\%AUC_{extrap}$ (%) | $1.86 \pm 0.844$ | $1.54 \pm 0.572$ |
| $C_{max}$ (ng/mL) | $6{,}350 \pm 978$ | $15{,}100 \pm 2{,}330$ |
| $t_{max}$ (h) | $1.03 \pm 0.0167$ | $1.06 \pm 0.0345$ |
| $t_{1/2}$ (h) | $4.12 \pm 0.458$ | $3.95 \pm 0.409$ |
| $DAUC_{0\text{-tlast}}$ (h•ng/mL/mg) | $412 \pm 70.2$ | $463 \pm 86.6$ |
| $DAUC_{0\text{-}\tau}$ (h•ng/mL/mg) | $412 \pm 70.4$ | $463 \pm 86.7$ |
| $DAUC_{0\text{-}\infty}$ (h•ng/mL/mg) | $420 \pm 74.9$ | $470 \pm 90.1$ |
| $DC_{max}$ (ng/mL/mg) | $99.3 \pm 15.3$ | $118 \pm 18.2$ |

**B**

| Parameter | **64 mg PA-001** (N = 5) | **128 mg PA-001** (N = 6) |
|---|---|---|
| $AUC_{0\text{-tlast}}$ (h•ng/mL) | $27{,}000 \pm 3{,}790$ | $62{,}300 \pm 13{,}300$ |
| $AUC_{0\text{-}\tau}$ (h•ng/mL) | $26{,}500 \pm 3{,}520$ | $61{,}200 \pm 12{,}700$ |
| $C_{max}$ (ng/mL) | $6{,}190 \pm 725$ | $16{,}100 \pm 2{,}370$ |
| $t_{max}$ (h) | $1.05 \pm 0.0391$ | $1.03 \pm 0.0139$ |
| $t_{1/2}$ (h) | $4.38 \pm 0.345$ | $4.54 \pm 0.771$ |
| $AR_{AUC}$ | $1.01 \pm 0.0606$ | $1.03 \pm 0.0556$ |
| $DAUC_{0\text{-tlast}}$ (h•ng/mL/mg) | $421 \pm 59.2$ | $486 \pm 104$ |
| $DAUC_{0\text{-}\tau}$ (h•ng/mL/mg) | $414 \pm 55.0$ | $478 \pm 98.9$ |
| $DC_{max}$ (ng/mL/mg) | $96.8 \pm 11.3$ | $126 \pm 18.5$ |

mean ± SD statistics presented

$AUC_{0\text{-tlast}}$: area under the curve from time 0 to the time of the last quantifiable concentration

$AUC_{0\text{-}\tau}$: area under the curve over a dosing interval ($\tau$, 24h)

$AUC_{0\text{-}\infty}$: area under the curve from time 0 extrapolated to infinity

$\%AUC_{extrap}$: percentage of area under the curve due to extrapolation from the last quantifiable concentration to infinity

$C_{max}$: maximum observed concentration

$t_{max}$: time of the maximum observed concentration

$t_{1/2}$: apparent terminal elimination half life

$DAUC_{0\text{-tlast}}$: $AUC_{0\text{-tlast}}$ normalized by dose administered

$DAUC_{0\text{-}\tau}$: $AUC_{0\text{-}24}$ normalized by dose administered

$DAUC_{0\text{-}\infty}$: $AUC_{0\text{-}\infty}$ normalized by dose administered

$DC_{max}$: $C_{max}$ normalized by dose administered

$AR_{AUC}$: accumulation ratio $_{0\text{-}\tau}$

**Figure Legend for Supplementary Figures**

**Supplementary Fig. 1. Schematic model for the conformational change of SARS-CoV-2 Spike protein during viral entry.** (a) SARS-CoV-2 Spike protein, comprising the S1 and S2 subunit, originally forms a locked trimer in the pre-fusion form. (b-c) By engagement of the host receptor, angiotensin converting enzyme 2 (ACE2), with the receptor-binding domain (RBD) of the S1 subunit, the S1-ACE2 complex dissociates from the S2 trimer. (d) The released S2 subunit undergoes the conformational change to form a long central coiled-coil that involves the extended HR1, with dynamic locational change of FP and the adjacent FPPR to form a blunt cone structure at the top and be inserted into the target cell membrane (intermediation form). (e-f) The coiled-coil structure subsequently refolds through the interaction of HR1 with HR2 and forms a six-helix bundle (post-fusion form), which bring viral envelope and cellular membrane closer together to fuse them. Anti-S1 (RBD) antibodies bind to RBD in S1 and inhibit the S1-ACE2 binding; anti-S2 antibodies and HR2-mimicking linear peptides bind S2 after the S1 release to inhibit the transition to the post-fusion form. In contrast, PA-001 binds to FPPR in the pre-fusion form and inhibits the conversion toward the post-fusion form.

**Supplementary Fig. 2. Dose-response curves for PA-001 and RDV against the individual SARS-CoV-2 clones (A8, B8, C8, D8, E8, F8, G8, and H8) or the wild type (WT).** (A) Individual dose-dependent curves shown in Fig. 2D-b for each clone. SARS-CoV-2 RNA in the culture supernatant of SARS-CoV-2 infected-VeroE6/TMPRSS2 cells upon treatment with various concentrations of (a) PA-001 or (b) RDV for 24 h was quantified and plotted against the drug concentration. (B) 90% inhibitory concentration ($IC_{90}$ in nM) of PA-001 and RDV against each clone was calculated by the linear regression of the dose-response curves shown in (A). (C) Substituted amino acids in the whole SARS-CoV-2 sequence in each clone, as compared to the original virus (WT).

**Supplementary Fig. 3. Anti-SARS-CoV-2 activity of PA-001 against Omicron sublineages.** (A) VeroE6/TMPRSS2 cells inoculated with SARS-CoV-2 pseudovirus carrying the Spike derived from Wuhan (a), JN.1 (b), or XFG (c) were treated with PA-001 or casirivimab + imdevimab (CAS + IMD) at various concentrations for 1 h. After additional culture without compounds or antibodies for 24 h, the cells were lysed and assessed for luciferase activity driven by SARS-CoV-2 pseudovirus infection. (B) Sequence alignment of aa 834 to 863 in the Spike region among the SARS-CoV2 Wuhan (WT), JN.1, XFG, and NB1.8.1. Asterisk in amino acids indicates full sequence conservation in all these variants.

#### Supplementary Fig. 1

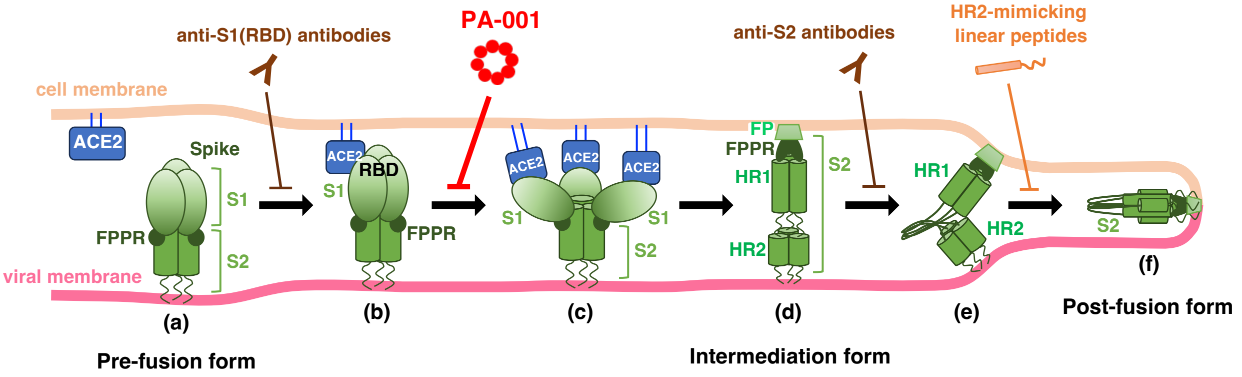

#### Supplementary Fig. 2

### A

a) Sensitivity to PA-001

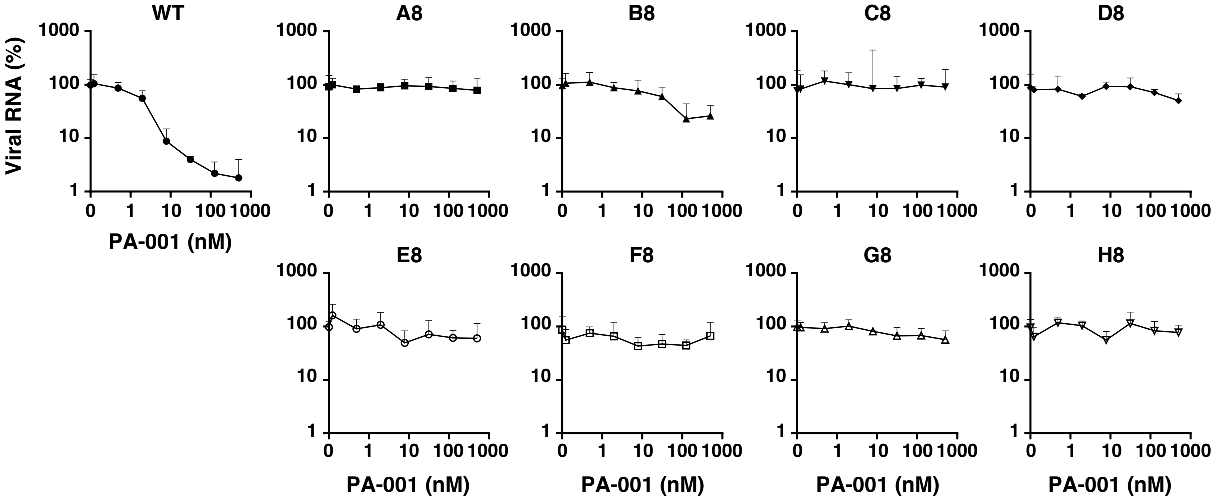


b) Sensitivity to RDV

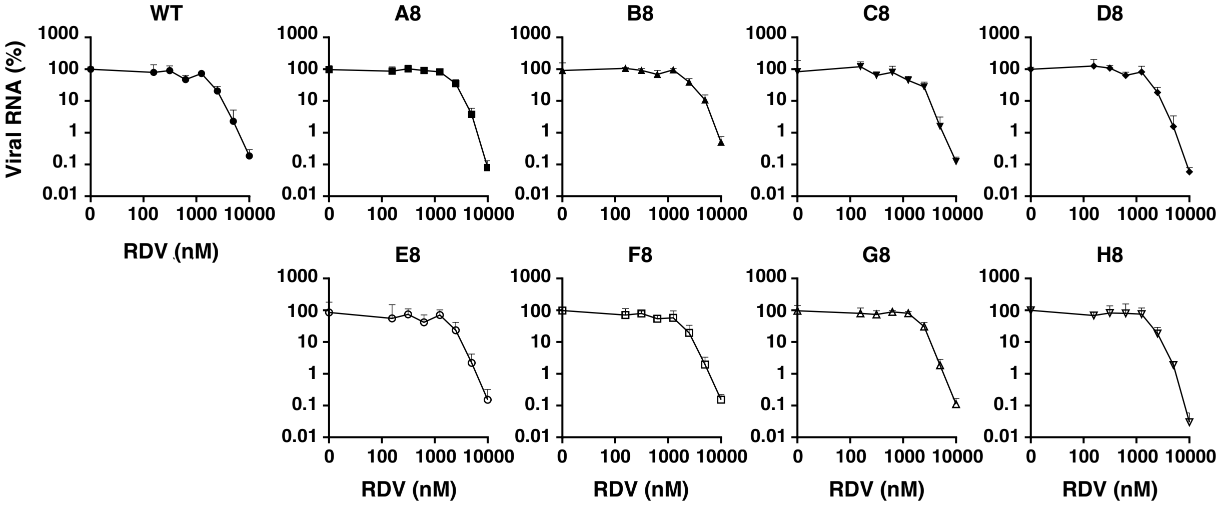


### B

**Sensitivity of each SARS-CoV-2 clone to PA-001 and RDV**

| Clones | $IC_{90}$ of PA-001 (nM) | $IC_{90}$ of RDV (nM) |
|---|---|---|
| WT | 8.39 | 3,310 |
| A8 | >500 | 3,850 |
| B8 | >500 | 4,480 |
| C8 | >500 | 3,000 |
| D8 | >500 | 3,180 |
| E8 | >500 | 3,320 |
| F8 | >500 | 3,080 |
| G8 | >500 | 3,340 |
| H8 | >500 | 3,140 |

### C

**Substituted amino acids in each SARS-CoV-2 clone**

| Clones | ORF1b | ORF7 | S1 | S2 |
|---|---|---|---|---|
| WT | - | - | - | - |
| A8 | - | Q94K | H655Y | Y837H |
| B8 | G662S | - | - | G838S |
| C8 | - | - | H655Y | L858I, N914Y |
| D8 | - | - | - | L841R |
| E8 | - | - | - | P862Q |
| F8 | - | - | - | Y837H, Q836del |
| G8 | - | - | - | P862Q |
| H8 | - | - | - | L841R |

### Supplementary Fig. 3

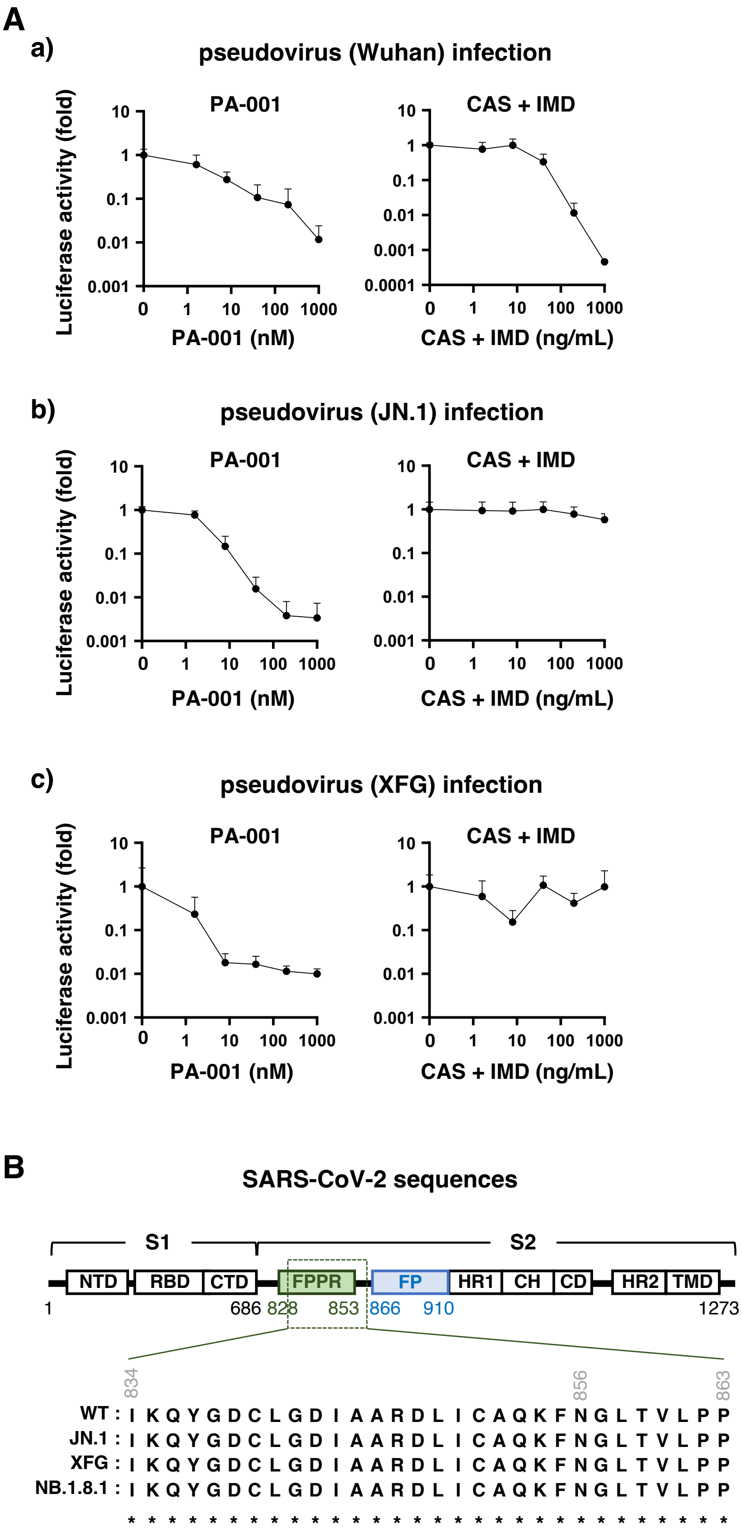
